# Direct cortical feedforward and feedback pathways differentially shape visual responses and selectively engage looped circuits

**DOI:** 10.64898/2026.10.27.762146

**Authors:** Flora Vasile, Matthijs N. Oude Lohuis, Camille Mazo, Devanshi P. Shah, Francesca Mastrogiuseppe, Christian K. Machens, Leopoldo Petreanu

## Abstract

Nearly every computational account of hierarchical cortex rests on assumptions about the influence of direct cortico-cortical inputs and which neurons they act upon. Yet, disentangling their direct functional contributions from indirect, multisynaptic influences remains difficult. We examined how feedforward (FF) and feedback (FB) pathways between mouse primary visual cortex (V1) and posteromedial cortex (PM) modulate visual responses, combining retrograde labelling, multi-area two-photon imaging and optogenetic perturbations. High or low activity in either pathway was associated with both excitation and inhibition of neurons in the targeted areas. FF activity was associated with stronger additive changes, while FB activity was related to stronger multiplicative modulations and improvements in visual decoding. Optogenetic manipulations causally confirmed these asymmetries. Both pathways preferentially modulated reciprocating “looped” neurons, with FF exciting them and FB either exciting or inhibiting them. These findings demonstrate that feedforward and feedback pathways serve distinct functional roles within recurrent cortical loops.

## Introduction

Visually-guided behavior and perception critically depend on neural activity distributed across a set of hierarchically organized visual areas of the neocortex. These areas are densely interconnected through a network of direct cortico-cortical connections that are anatomically asymmetric: feedforward (FF) and feedback (FB) inputs originate from neurons in different cortical layers and innervate distinct layers within their target areas^1–4^. They also innervate separate dendritic compartments of pyramidal neurons^5,6^ and differentially target interneuron subtypes^4,7–9^. Functionally, FF inputs are thought to drive activity in their target areas, whereas FB inputs are generally considered modulatory. However, the evidence for this functional asymmetry is mixed. In brain slice preparations, FF and FB inputs have been shown to act both as driver and modulators^10^. In vivo, FB inputs appear capable of having driving, additive effects on their target regions in the absence of feedforward visual input, such as in response to occluded images^11,12^ or stimuli that elicit the percept of illusory contours^13,14^. Moreover, the modulatory effects attributed to FB are themselves diverse, including multiplicative gain modulation, such as when mediating attentional effects^15,16^, and subtractive influences when cancelling expected stimuli^17,18^.

Clarifying the influence of FF and FB interactions requires isolating the contributions of direct cortico-cortical inputs from those of other pathways that may also influence activity in the target areas. However, manipulations of neural activity of an area, either by cooling^19,20^, pharmacological silencing^21^, electrical microstimulation^22^ or optogenetic manipulations^23–26^ while measuring the consequences in another cannot distinguish whether the observed effects are mediated by direct projections from the manipulated area to the target or by alternative pathways, such as those involving higher-order thalamic nuclei^27,28^. In the few cases where cortico-cortical pathways have been selectively manipulated, this has typically been done in only one direction^29–31^, precluding a direct comparison of the differential influences of FF and FB inputs. Similarly, methods leveraging statistical relations in simultaneous multi-area recordings to infer inter-area interactions^32–35^ cannot determine whether the inferred interactions arise from direct cortico-cortical connections or from indirect pathways.

Beyond distinguishing FF and FB influences in general, cortico-cortical inputs might exert different influences depending on the projection type of the target neuron. For example, predictive coding requires FB to selectively exert subtractive influences in neurons harboring predicted representations that send reciprocating projections to the source of the FB inputs^17,18^. While there is anatomical^36^ and circuit mapping evidence^6,37^ showing that cortico-cortical projections might differentially engage with reciprocating, looped neurons, i.e. neurons that project back to the source, whether FF and FB pathways exert distinct influences on reciprocating versus non-reciprocating neurons during visual processing remains unknown.

To fill these gaps, we combined statistical methods for inferring the influence of direct cortico-cortical inputs with retrograde labeling and pathway-specific optogenetic manipulations to determine: (1) how direct feedforward and feedback inputs shape visual representations, and (2) whether these pathways exert distinct influences on reciprocally connected neurons versus neurons that do not project back to the source region. We found that FF and FB pathways both exert mixed excitatory and inhibitory effects, but FF activation preferentially drives additive excitation in PM responses, whereas FB produces stronger multiplicative gain modulation in V1 and greater improvements in visual decoding. Both pathways preferentially engage looped neurons, with FF more strongly favoring their excitation, whereas FB is more likely to excite and inhibit them. Our results identify distinct, asymmetric roles for the two pathways in shaping visual representations and engaging recurrent hierarchical computations.

## Results

### Measuring the influence of direct cortico-cortical inputs in multi-area recordings

To measure the influence of cortico-cortical projections on visual representations, we performed simultaneous calcium imaging in two visual areas while identifying neurons that send projections between them. Using a combination of retrograde adeno-associated viruses (AAVs) expressing Cre and Flpo recombinases with others encoding for Cre- and Flp-dependent tdTomato expression, we labeled V1 neurons projecting to PM (V1_PM_; FF-projecting neurons) as well as PM neurons projecting to V1 (PM_V1_; FB-projecting neurons) within the same GCaMP6s-expressing transgenic mice (Fig. 1A,B). FF-projecting cells in V1 were predominantly found in superficial layer 2/3, whereas FB projecting cells were more widely distributed across layers and most numerous in deeper layers (Fig. 1C). Using a two-photon random access large field of view mesoscope^38^ we simultaneously recorded from locations in V1 and PM with tdTomato-expressing neurons (Fig. 1D) as mice were presented with moving gratings while awake over a treadmill (n = 12 sessions from N = 8 mice). On average, we identified 113.3±71.0 labeled cells per session, corresponding to 16.4±16.9 labeled cells per imaging plane in V1 (range: 0-77) and 13.9±16.4 in PM (range: 0-64).

**Figure 1:**
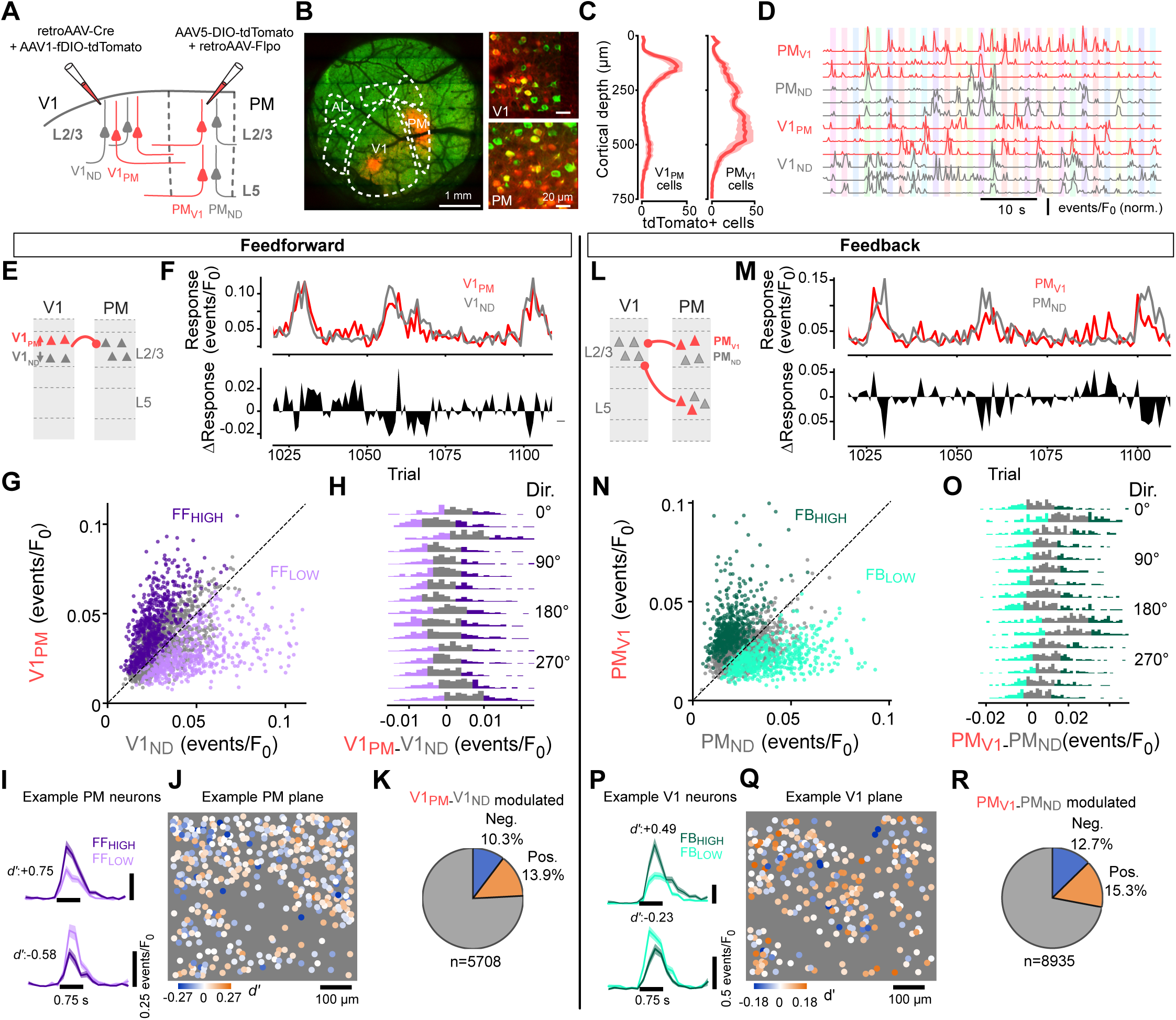
Relating cortico-cortical pathway activity to visual responses in multi area recordings. **A)** Viral approach to label PM-projecting V1 neurons (V1PM) through retrograde Flpo-mediated viral expression and V1-projecting PM neurons (PMV1) through Cre-mediated expression. **B)** Example cranial window with GCaMP6s (green) and tdTomato (red) fluorescence and insets showing tdTomato-labeled and unlabeled cells. White dotted lines indicate areal borders determined with intrinsic imaging. **C)** Mean number of labeled FF-projecting and FB-projecting cells per imaging plane as a function of cortical depth ( N = 12 mice). Shading, s.e.m. **D)** Example traces of simultaneously imaged reciprocally projecting neurons in V1 and PM during drifting grating stimulation. Colored shading, stimulus period. **E)** Schematic approach to study the effect of endogenous FF activity on Layer 2/3 of PM. The difference in activity between the V1PM and V1ND populations was taken as a proxy of endogenous fluctuations in FF activity to PM while controlling for overall activity fluctuations in V1. **F)** Top, mean response of the V1PM and V1ND populations for several trials in an example session. Bottom, difference between the two. **G)** Mean activity of the V1PM and V1ND populations across all trials for an example session (n = 2234 trials). Trials classified as FFLOW or FFHIGH are colored. **H)** Each row is a histogram of V1PM–V1ND activity across trials of the same grating direction for the example session. FFLOW and FFHIGH states were identified per stimulus direction as the 25% trials with highest and lowest activity difference. **I)** Activity of two example neurons in PM whose response was significantly positively or negatively modulated by the state of the feedforward pathway (*d’* of FFLOW vs. FFHIGH). **J)** Example imaging plane in L2/3 with PM cells color coded by the modulation strength. *d’*, discriminability index. **K)** Fraction of PM neurons negatively (blue) and positively (orange) modulated by FF activity. **L-M**) Same as in panels E-F but for the PMV1 and PMND populations in the same session. **N-O**) Same as in panels G-K but for the relation of the FB-projecting population of PM on V1 activity. Same example recording session as in G-K.

To measure the influence of cortico-cortical projections on their target areas we focused on endogenous fluctuations in the level of FF or FB activity and identified periods in which the average activity of the V1_PM_ or PM_V1_ neurons was either high or low. To avoid confounding these periods with global cortical modulations driven by behavioral state^39–41^ we focused our analyses on trials without locomotion and discernable facial motion. However, even during these still trials mean activity in V1 and PM was positively correlated (r = 0.44±0.17, n = 12 sessions, p = 7.21 x 10^-6^, one sample t-test). Therefore, to control for global rate fluctuations shared across the mouse visual system, we compared activity levels between labeled and neighboring unlabeled neurons in the source area. Specifically, periods of low or high FF activity were identified by the difference in average activity between V1_PM_ neurons and nearby unlabeled neurons (V1_ND_) (Fig. 1E,F). This metric was not significantly correlated with PM activity across sessions (correlation of V1_PM_ – V1_ND_ activity with PM L2/3: r = 0.11±0.17, p = 0.07, one sample t-test, n = 10 sessions). For each stimulus, we identified trials with high FF activity (FF_HIGH_) as the top 25% of trials where V1_PM_ neurons were most active relative to V1_ND_ neurons. Conversely, periods of low FF activity (FF_LOW_) were defined as the 25% of trials showing the smallest activity difference between the two populations (Fig. 1G,H). This approach yielded an equal number of FF_HIGH_ and FF_LOW_ trials for each stimulus, ensuring that the FF_HIGH_ and FF_LOW_ do not simply reflect potential direction tuning differences between the imaged V1_PM_ and V1_ND_ populations. FF_HIGH_ trials resulted in either positive or negative modulation of PM neuronal visual responses relative to those observed during FF_LOW_ trials (Fig. 1I). Positively and negatively modulated neurons were found in approximately equal proportions (Fig. 1J, K). While excited and inhibited cells were intermingled, modulation strength was more similar between nearby neurons than between distant ones (Fig. S1A).

Using the same approach comparing the activity of PM_V1_ with neighboring unlabeled neurons (PM_ND_) neurons, we identified trials with high or low FB activity (FB_HIGH_ and FB_LOW_) (Fig. 1L,M). As with FF, the difference in activity between the two projection-defined populations was not correlated with V1 activity (correlation of PM_V1_ – PM_ND_ with V1 L2/3: r = 0.08±0.17, p = 0.16, one sample t-test, n = 12 sessions) nor with the FF activity metric (correlation of PM_V1_ – PM_ND_ with V1_PM_ – V1_ND_ activity: r = 0.01±0.12 across sessions, p = 0.79 one sample t-test, n = 10 sessions). V1 neurons were positively and negatively modulated during FB_HIGH_ periods in similar proportions (Fig. 1 P-R). As with FF inputs, nearby neurons exhibited more similar modulation by FB activity than neurons located farther apart (Fig. S1A).

We tested whether the modulations observed in V1 and PM, as a function of FF- and FB-projecting neuron activity, reflect direct influences of these neurons by additionally recording from the anterolateral visual area (AL) in a subset of experiments. If the modulation of PM neurons by FF_HIGH_ and FF_LOW_ states in V1 reflects direct cortico-cortical interactions, modulations should be weaker in AL than in PM. Consistent with this prediction, a smaller fraction of neurons in AL exhibited significant modulation of their visual responses by FF_HIGH_ and FF_LOW_ compared with area PM (Fig. S2A). Similarly, fewer neurons in area AL were modulated by FB_HIGH_ and FB_LOW_ than in V1, consistent with these modulations reflecting direct influences of FB projections to V1 (Fig. S2B).

### Asymmetric influences of FF and FB pathways on visual responses

We next investigated whether neuronal tuning to drifting gratings depended on FF or FB activity by comparing responses to different grating directions in one region during trials when FF or FB activity was high or low in the other region. The preferred direction or tuning width did not vary with FF or FB activity (Fig. S3). However, fluctuations in population activity can modulate the tuning of V1 neurons additively and multiplicatively while leaving tuning unchanged (Fig. 2A,C)^42,43^. To assess the extent of additive and multiplicative influences in cortico-cortical interactions, we fitted a linear regression to the average responses to the different grating directions during FF_LOW_ and FF_HIGH_ trials for positively modulated PM neurons (Fig. 2A,B), and during FB_LOW_ and FB_HIGH_ trials for positively modulated V1 neurons (Fig. 2C,D). The offset of these linear fits captures additive or subtractive shifts of the tuning curve while the slope captures multiplicative or divisive effects. We assessed the significance of both terms with a permutation test. When focusing on positively modulated V1 neurons, the average slope of these linear fits was greater than 1, indicating a predominantly multiplicative effect of FB activity. The slope describing FF influences on positively modulated PM neurons was closer to 1 on average and significantly lower than that measured for FB interactions (FF: 0.903, FB: 1.523, p = 2.35 x 10^-49^, Mann-Whitney U test). In contrast, the average additive offset was larger for FF influences when compared to the FB ones (Fig. 2E) (FF: 0.015 events/F_0_, FB: 0.006 events/F_0_, p = 1.73 x 10^-29^, Mann-Whitney U test).

**Figure 2:**
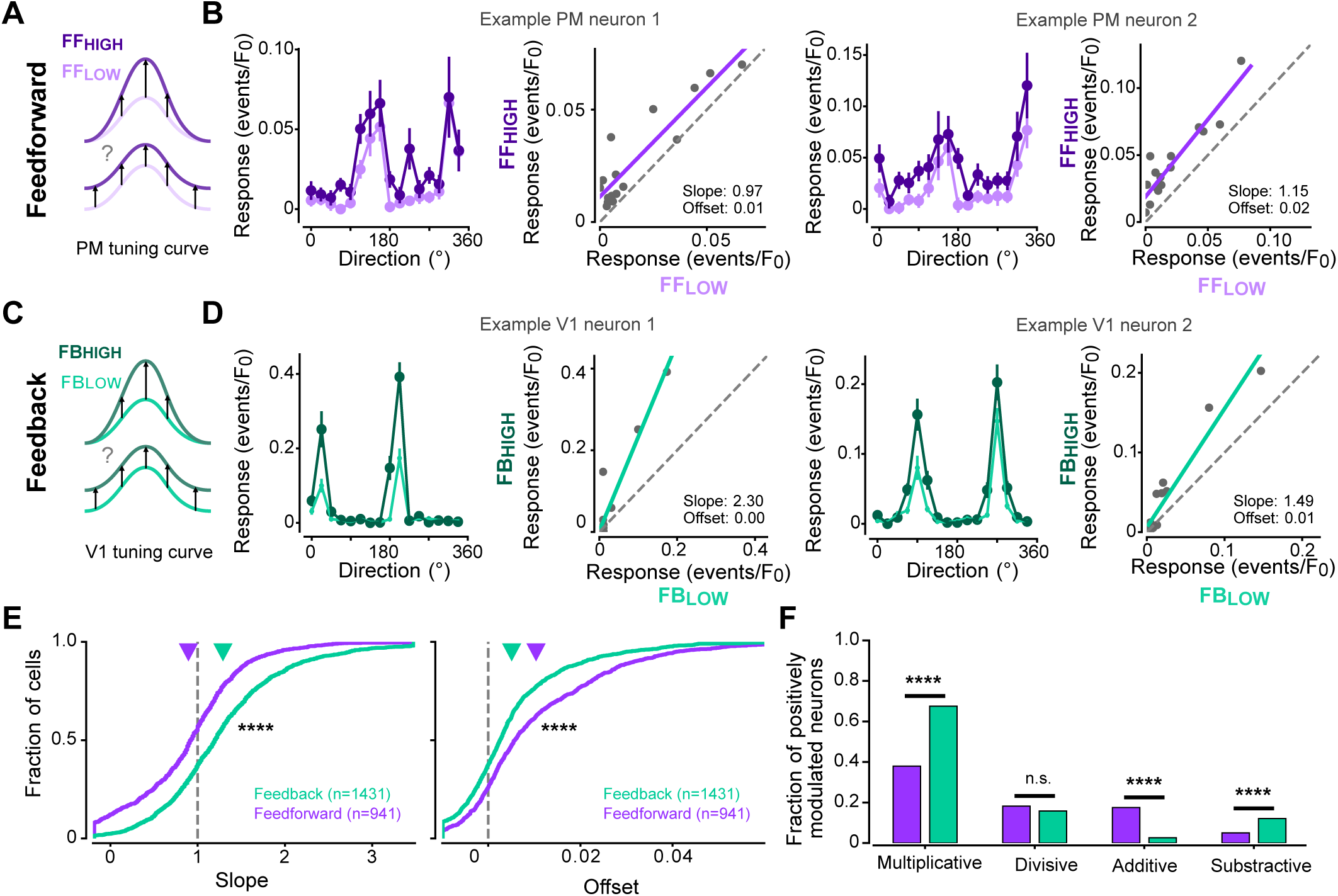
FF exerts more additive, whereas FB exerts more multiplicative, influences on visual representations. **A)** Cartoon on how tuning curves would differ in PM neurons between FF_HIGH_ and FFLOW states in the case of a pure multiplicative (top) or additive (bottom) influence. **B)** Tuning curves in two example PM neurons during FF_HIGH_ and FFLOW states. Left, mean response to each stimulus direction. Errorbars, s.e.m. across trials. Right, same data as a scatter plot with regression fit indicating a predominantly additive effect of FF activity. **C)** Cartoon of multiplicative of additive modulation by the FB pathway on V1 tuning curves. **D)** Tuning curves and regression fits for two example V1 neurons showing multiplicative modulation by FB activity. **E)** Cumulative distribution of slope (left) and offset (right) of all positively correlated cells. Upper triangles indicate the mean. Neurons sampled during 12 sessions across N=8 mice. Here and throughout all figures: *p<0.05, **p<0.01, ***p<0.001, ****p<0.0001. **F)** Fraction of cells (out of all excited cells) that show significant gain or offset modulation.

Consistent with the measurements of slope and offset, the fraction of significantly multiplicatively modulated neurons was higher for FB influences, whereas additively modulated neurons were more prevalent for FF influences (Fig. 2F) (multiplicative: χ^2^(1,2372) = 130.61, p = 2.4 x 10^-29^; additive: χ^2^(1,2372) = 112.15, p = 2.7 x 10^-25^; χ^2^ test with Bonferroni correction). Subtractive modulation was also more common for FB than FF influences (χ^2^(1,2131) = 18.68, p = 0.00012). The fraction of neurons showing divisive effects was similar across both conditions (χ^2^(1,2131) = 4.12, p = 0.34). We did not find differences in the average slope and offset between the pathways for target neurons that were negatively modulated by FF and FB activity (Fig. S4A). Inhibited neurons consisted mostly of a divisive population with only small differences between pathways (Fig. S4B). The differences in multiplicative and additive modulation between FF and FB influences were not explained by differences in tuning or activity levels between PM and V1 neurons, as they persisted after controlling for mean evoked activity, response range, and orientation selectivity (Fig. S5A-C).

### Optogenetic manipulation confirms asymmetric influences of FF and FB pathways

The preceding analyses of multi-area recordings suggest that both FF and FB pathways can elicit positive and negative modulations of visual responses in their target areas. They further indicate that positive modulations driven by FF projections are relatively more additive than those elicited by FB inputs, whereas positive modulations driven by FB inputs are comparatively more multiplicative. However, statistical methods cannot establish causality, and the observed covariances may be driven by shared inputs rather than direct interactions between neuronal populations^34,44^. To address these issues, we complemented the multi-area recordings with projection-specific optogenetic manipulations between V1 and PM to establish the causal contributions of FF and FB pathways to the modulation of visual responses in their target areas.

To measure FF influences, we injected AAVs expressing the red-shifted opsin ChrimsonR^45^ into V1 in GCaMP6-expressing mice and recorded visual responses in PM (Fig. 3A). Conversely, to study FB influences, we injected the virus in PM and recorded in V1. In both cases we included in the injection pipette retro-AAVs^46^ resulting in mCherry expression in neurons projecting back to injection site. Because FF and FB projections in visual cortex are retinotopically matched^47,48^ and the influence of cortico-cortical inputs depends on the retinotopic distance between projecting axons and recorded neurons^49^ this approach allowed us to consistently position our two-photon recordings in equivalent retinotopically matched locations by centering the recording over the retrogradely labelled mCherry expressing neurons (Fig. 3B). In addition, it allowed comparing the influence of FF and FB pathways on the reciprocating “looped” neurons (PM_V1_ and V1_PM,_ respectively) and neurons with undetermined projection patterns (PM_ND_ and V1_ND,_ respectively). Injections in V1 resulted in ChrimsonR-expressing axons spanning most of the cortical depth but with higher density in the supragranular layers (Fig. 3B,C). In contrast, injections in PM produced ChrimsonR-expressing FB axons in V1 with a complementary laminar pattern, peaking in layer 1. As before, mCherry-expressing, FF-projecting V1_PM_ neurons were mostly in upper L2/3, while FB-projecting PM_V1_ were concentrated mainly in lower L2/3 and L5 (Fig. 3B and Fig. S7A-D). We identified an average number of looped cells per session of 16.2±10.3 in V1 (range: 3-42) and 8.4±6.6 in PM (range: 0-31). To measure the influence of FF and FB projections we photostimulated ChrimsonR-expressing axons through the imaging objective using a red laser as mice viewed moving gratings (Fig. 3D). Photostimulation of FF or FB axons during sensory stimulation elicited both excitation and inhibition across neurons (Fig. 3D-J). As observed in the multi-area recordings, excited and inhibited neurons were intermingled, yet modulation strength was more similar among spatially proximal neurons than among more distant ones (Fig. S1B). These modulations were absent in mice that did not express ChrimsonR in cortico-cortical axons (Fig. S6), confirming they were not caused by non-specific effects of the laser light due to propagation through brain tissue^50^.

**Figure 3.**
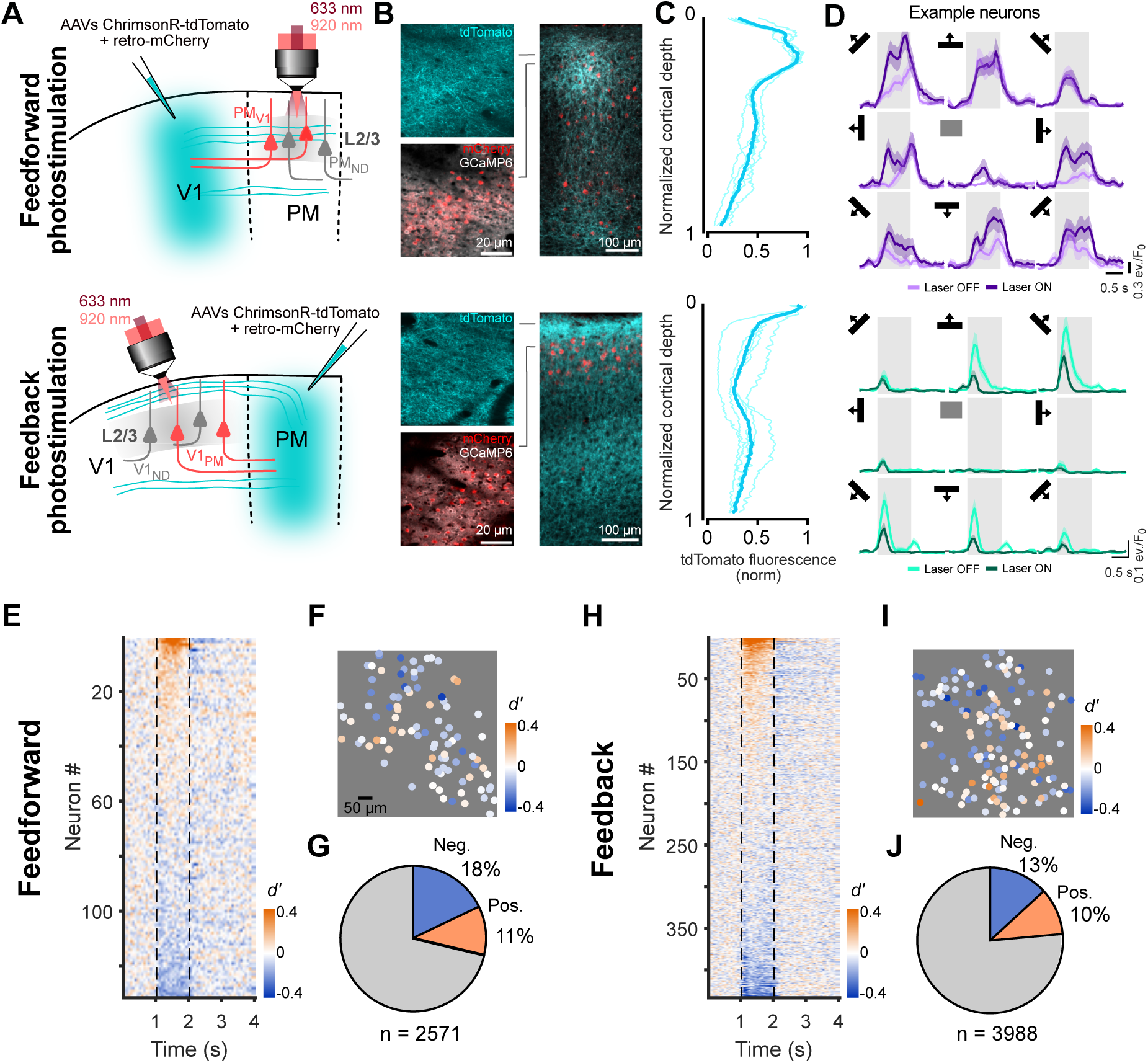
Optogenetic photostimulation of FF and FB pathways. **A)** Experimental approach. Top: FF V1PM axon photostimulation and retrograde labelling of PMV1 FB-projecting cells. Bottom: FB PMV1 axon photostimulation and retrograde labelling of V1PM FF-projecting cells. Axons were photostimulated through the imaging objective. **B)** Left: example field of view from two-photon recording. ChrimsonR-tdTomato-expressing axons (cyan), mCherry retrograde labelled cells (red) and GCaMP6f (grey). Right: Coronal section showing the laminar distribution of ChrimsonR-tdTomato-expressing axons and mCherry retrograde labelled cells. **C)** Laminar distribution of ChrimsonR-tdTomato-expressing FF axons in PM. Individual mice, thin traces; average, thick trace. (n= 5 mice). **D)** Example responses to gratings moving in different directions with and without photostimulation of FF (top) and FB (bottom) axons. Center trace, photostimulation in absence of gratings. Grey shading, stimulus and photostimulation period. **E)** Average discriminability index between Laser ON and Laser OFF per frame for all visually responsive neurons in PM from an example mouse with FF photostimulation. Dotted line, stimulus period. Orange and blue colors represents positive and negative modulations, respectively. **F)** Example field of view representing the neurons’ modulation. **G)** Fraction of significantly positively (orange) or negatively (blue) modulated visually responsive neurons. **H-J**) Same as E-G for FB photostimulation.

We compared the tuning curves of PM and V1 neurons with and without concurrent photostimulation of V1_PM_ or PM_V1_ axons, respectively (Fig. 4A,B). Photostimulation of FF or FB axons did not change the preferred orientation or tuning width of the neurons in their target areas (Fig. S3E-H), consistent with the multiarea recordings. We compared mean neuronal responses to each moving grating in trials with and without photostimulation, then used linear regression to quantify the additive and multiplicative changes induced by activating FF or FB axons, as in the multiarea recordings. In neurons that were excited by photostimulation, activation of V1_PM_ axons resulted in a smaller slope in PM responses than activation of PM_V1_ axons did in V1 (Fig. 4C; FF: 0.94±0.48, FB: 1.38±0.82, p = 9.9 x 10^-6^, Mann-Whitney U test, FF: n = 66 neurons from N = 13 mice, FB: n = 95 neurons from N = 9 mice). In contrast, the opposite pattern was observed for the additive effect of photostimulation: activation of V1_PM_ axons produced a larger additive offset in PM responses than activation of PM_V1_ axons did in V1 (FF: 0.06±0.14 events/F_0_, FB: 0.03±0.07 events/F_0_, p = 0.008, Mann-Whitney U test). Consistent with this, although photostimulation of both FF and FB pathways produced additive and multiplicative effects in a large fraction of neurons, a greater fraction of neurons showed multiplicative modulation following FB stimulation than following FF activation (odds ratio = 2.8, p = 0.002, Fisher’s exact test) (Fig. 4D). No differences were found in the offset or slope in negatively modulated neurons (Fig. S4C-D). The differences in slope and offset when photostimulating FF and FB axons could not be attributed to differences in the amplitude or range of the visual responses or the magnitude of the modulatory effects in V1 and PM neurons (Fig. S5 D-F). We conclude that the pathway-specific optogenetic manipulations causally confirm the observations from the multiarea recordings, showing that FF and FB projections exert distinct excitatory influences: FF inputs are more additive, whereas FB inputs have a more multiplicative effect in their respective target areas.

**Figure 4.**
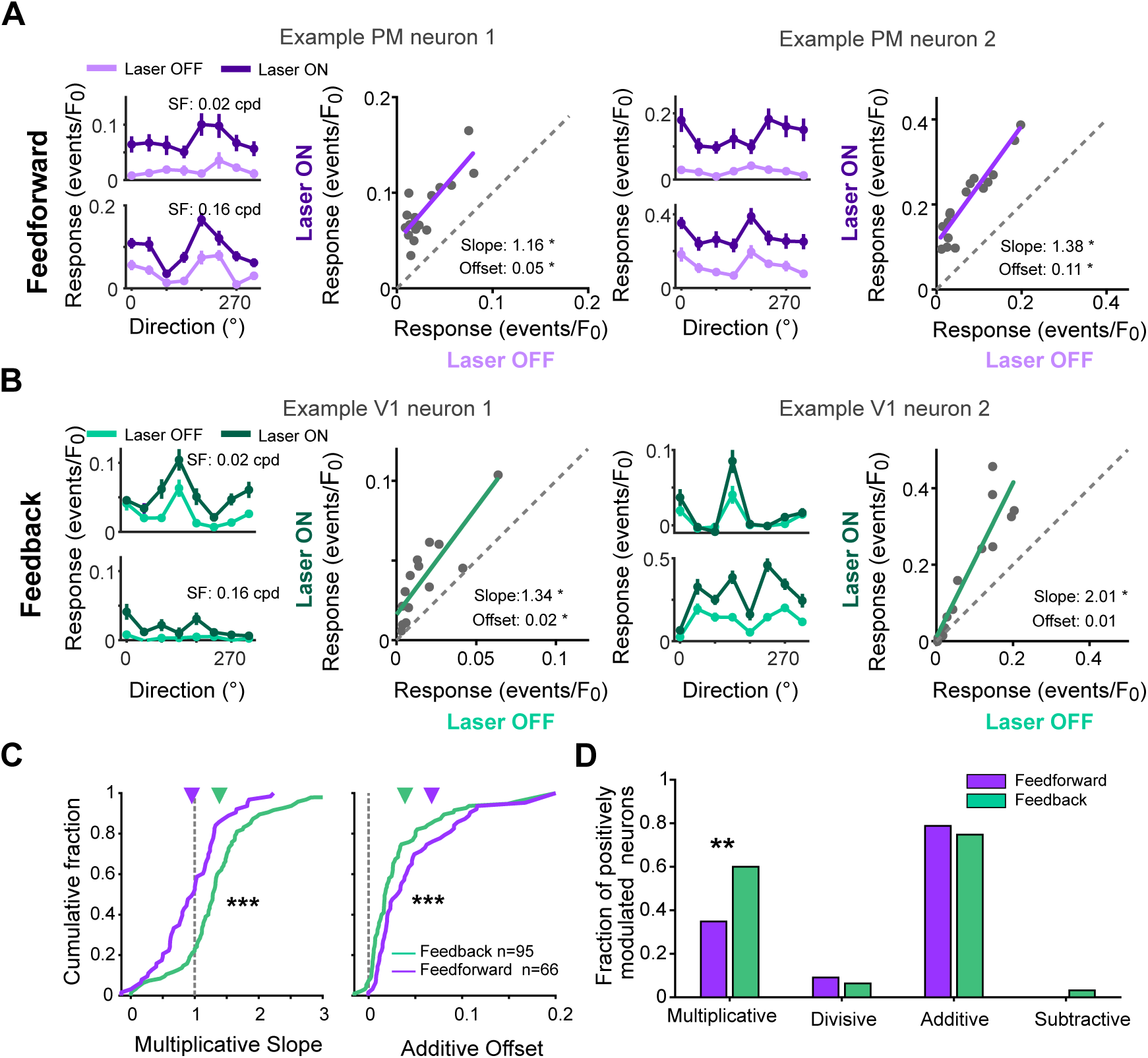
Photostimulation of FF and FB pathways induce different modulations. **A)** Tuning curves and regression fit for two example excited PM neurons upon FF photostimulation. Left: neuronal responses to gratings moving in different directions with Laser OFF or Laser ON. Top and bottom plots show responses of the same neuron to gratings of different spatial frequencies (SF). Right: regression fit of the responses during Laser OFF vs. Laser ON trials for gratings at the two spatial frequencies and photostimulation without visual stimulus. **B)** Same as A for two example V1 neurons during FB photostimulation. **C)** Cumulative distribution of the slope (left) and offset (right) of cells excited by FF or FB photostimulation. Neurons were sampled from : FF, 30 sessions from N=13 mice; FB, 20 sessions from N=9 mice. **D)** Fraction of neurons with significant multiplicative, divisive, additive and subtractive modulations during FF and FB photostimulation.

### Feedforward and feedback cortico-cortical inputs differentially affect stimulus representations

Multiplicative and additive effects can affect encoding of sensory information differentially^42,43,51^. Thus, we investigated how the differences in modulatory influence on excited neurons (additive versus multiplicative) between the two types of corticocortical pathways affect visual representations. We estimated the quality of representations by training linear readouts to decode stimulus identity from neural activity. While decoding performance from V1 activity was higher in FB_HIGH_ than in FB_LOW_ trials, performance in PM did not vary with the level of FF activity (Fig. 5A,B, FF: p = 0.84, paired t-test, n = 10 sessions; FB: p = 0.0032, n = 12 sessions). This finding was replicated using decoders trained on recordings from the optogenetic experiments: decoding performance increased when FB axons were stimulated, but not when FF axons were stimulated (Fig. 5D,E, FF: p = 0.57, FB: p = 0.004, paired t-test). The improvements in decoding performance under FF_HIGH_ were due to multiplicative gain changes, as they depended on the average slope but not the average offset of the randomly selected neuronal populations used for decoding (Fig. 5C,F; multiarea recordings: correlation between decoding improvement and mean slope, FF: r = -0.03, p = 0.28, FB: r = 0.47, p = 2.1 x 10^-30^; offset, FF: r = 0.04, p = 0.24; FB: r = -0.16, p = 3.2 x 10^-9^, optogenetic experiments: slope, FF: r = 0.09, p = 0.03, FB: r = 0.23, p = 1.9 x 10^-7^; offset, FF: r = 0.007, p = 0.88; FB: r = -0.13, p = 2.7 x 10^-3^). Thus, the predominantly multiplicative modulation of visual responses exerted by FB inputs enhances stimulus discriminability, whereas the comparatively more additive modulation mediated by FF drive does not.

**Figure 5:**
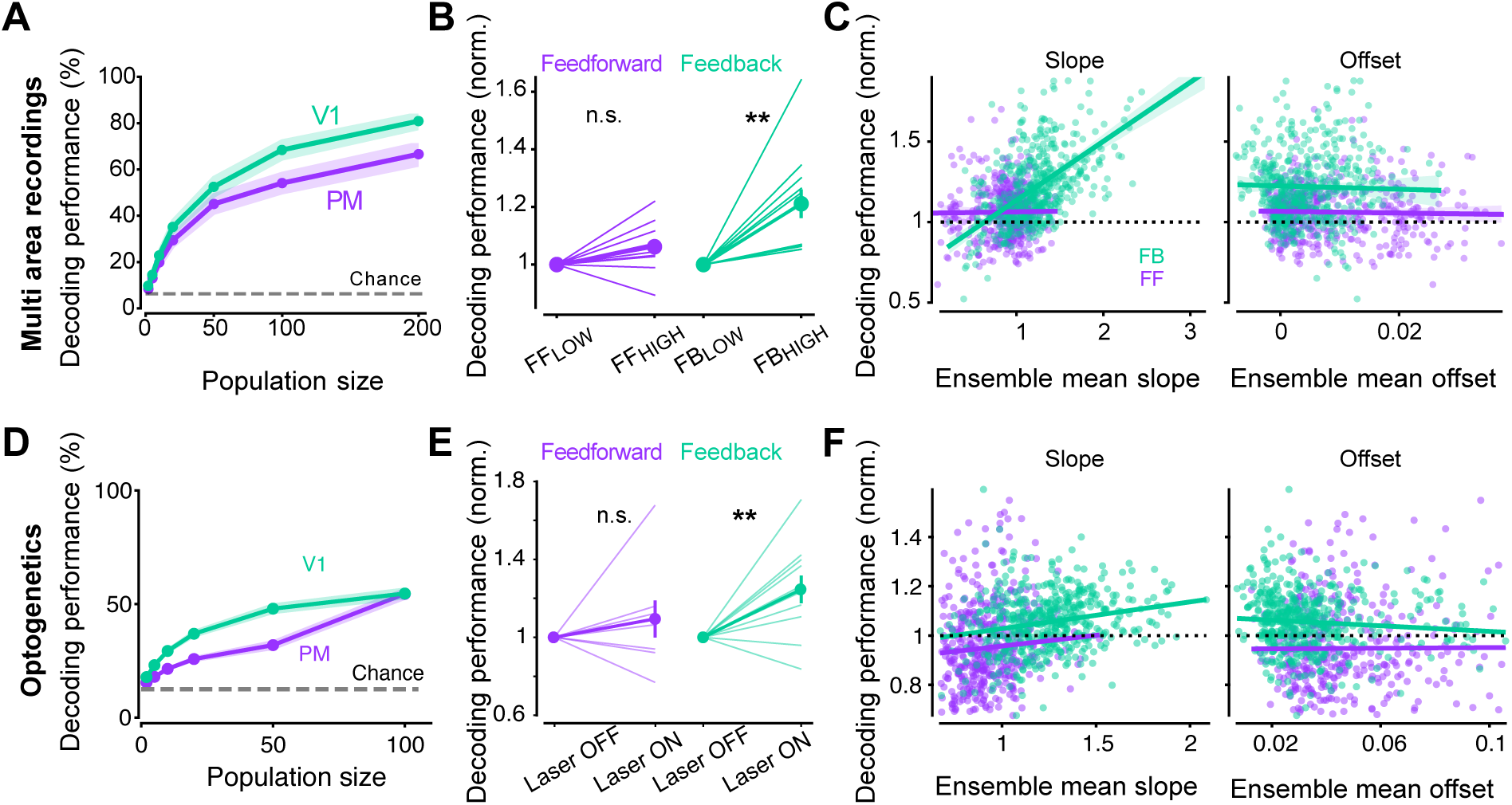
Increased feedback activity improves visual stimulus decoding through multiplicative gain modulation. **A)** Cross-validated decoding performance with increasing size of population of PM and V1 neurons used for decoding stimulus direction in multi area recordings. Line, mean; shading, s.e.m. across sessions (n=12 from N=8 mice). Dotted line, chance level. **B)** Performance of decoding from trials with low or high FF or FB activity in multi-area recordings. Values are normalized to the performance on low trials per session. Mean, s.e.m. FF, n=10 sessions. FB, n=12 sessions from N=8 mice. **C)** Performance as a function of the average slope (left) and offset (right) of the neurons used for decoding (n=600 sampled ensembles of 10 neurons for each pathway from n=10/12 FF/FB sessions from N=8 mice). **D-F**) Same as A-C. for optogenetics experiments. **D)** Line, mean; shading, s.e.m. across sessions (FF: n=43 from N=16 mice, FB: n=27 from N=12 mice). Dotted line, chance level. **E)** Mean, s.e.m across sessions. FF: n=8 from N=5 mice, FB: n=11 from N=7 mice. **F)** n=500 sampled ensembles of 10 neurons for each pathway; FF: n=77 neurons, FB: n=113 neurons.

### Feedforward and feedback cortico-cortical input selectively modulate reciprocating neurons

We then tested whether photostimulation of FF and FB projections modulated neurons differently according to their projection type by comparing responses in neurons projecting back to the source area of the input (“looped” cells) with those of neurons with undetermined projection patterns. To do so, we compared the photostimulation-induced modulation of retrogradely labeled, mCherry-expressing neurons with that of nearby unlabeled neurons, thereby controlling for spatial heterogeneity in photostimulation effects (See Methods). We compared the fraction of looped neurons that were excited, inhibited, or modulated (excited or inhibited combined) during visual stimulation with the corresponding fraction in resampled ensembles of non-looped neurons, generated by randomly sampling one unlabeled neuron within 50 µm of each looped neuron (Fig. 6A). The photostimulation of FF-projecting V1_PM_ axons was more likely to result in a positive modulation of the visual responses in looped PM_V1_ neurons than in nearby non-looped neurons (p = 0.01, two-tailed resampling test) (Fig. 6B). The fraction inhibited was similar in both projection types, as well as the fraction modulated (pooling the fraction excited and inhibited). Similarly, photoexcitation of FF axons was more likely to recruit looped PM_V1_ neurons in the absence of visual stimulation (p = 0.005, two-tailed resampling test) (Fig. 6C). The selective excitation of looped neurons was robust across different distance thresholds used to select the non-looped neurons neighboring of PM_V1_ neurons (Fig. S7I,J). Using immunohistochemical analyses, we confirmed that retrogradely labeled mCherry-expressing neurons did not express the opsin, ruling out the possibility that the increased probability of looped PM neurons being excited by FF photostimulation was due to these neurons being directly photosensitive (see Methods).

**Figure 6:**
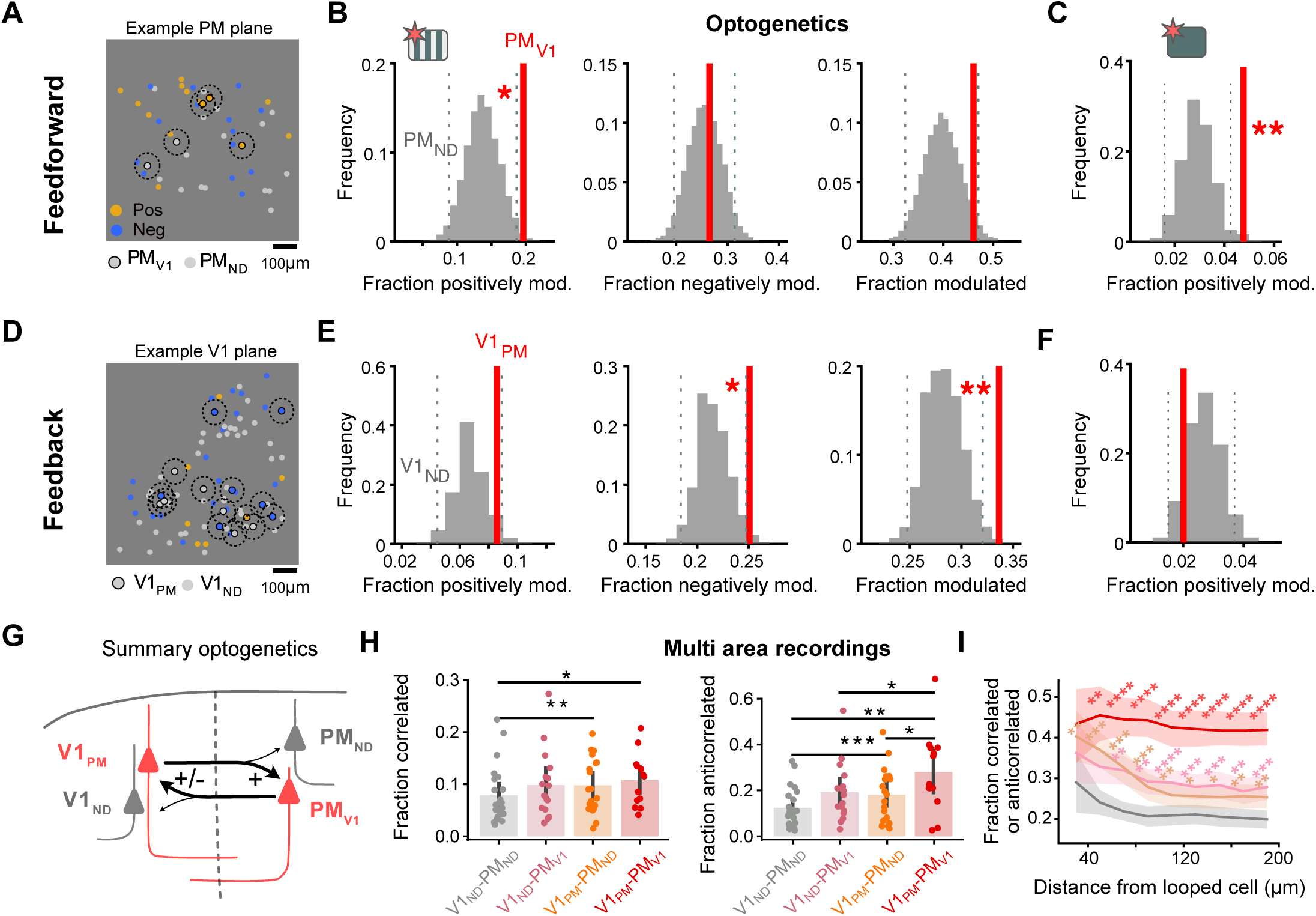
Selective and asymmetric modulation of reciprocally projecting neurons by FF and FB inputs. **A)** Example two-photon field of view of PM depicting the spatial distribution of neurons that are either excited (orange) or inhibited (blue) by FF photostimulation. Dotted circles depict the area within 50 μm of PMV1 (looped) neurons from which PMND neurons are selected for further analyses. **B)** Fraction of excited, inhibited and modulated PMV1 (red line) and shuffled resamples from PMND neurons located <50 μm from a PMV1 neuron (grey distribution, 10.000 resamples). See Figure S6 for results with other distance thresholds. n = 102 visually responsive PMV1 neurons and n = 310 PMND neurons from N = 15 mice. **C)** Same as B for the fraction of excited cells by photostimulation without visual stimulus. n = 377 PMV1 neurons and n = 1715 PMND neurons from N=18 mice. **D-F**) Same as A-C for the effects of FB photostimulation on V1 neurons. E, n=315 V1PM neurons and n=1164 V1ND neurons from N = 17 mice. F, n=595 PMV1 neurons and n = 2587 PMND neurons from N = 17 mice. **G)** Summary of results. Photostimulating FF V1PM axons preferentially excites PMV1 neurons and photostimulating FB PMV1 axons selectively modulates (excite or inhibit) V1PM neurons. **H)** Fraction of neuron pairs that is significantly correlated (left) or anticorrelated (right) for different population pairs. Asterisks denote significance of a paired t-test with Bonferroni multiple comparison correction. n=13 sessions from N = 8 mice. Only non-looped neurons within 50 μm distance of a looped cell were considered. **I)** Fraction of neuron pairs that is either correlated or anticorrelated in the multi area recordings while varying the distance threshold to compare neighbouring labelled and unlabelled neurons. Line, mean; shading s.e.m across sessions. Asterisks denote significance tested against V1ND-PMND pairs at each distance threshold, paired t-test with Bonferroni multiple comparison correction.

Photostimulation of the FB axons of the PM_V1_ neurons also resulted in the selective modulation of looped neurons in V1 (V1_PM_). However, in contrast to FF photostimulation, this manipulation resulted in a larger fraction of looped neurons showing modulation (either excitation or inhibition) compared to their nearby non-looped neurons (p = 0.002, two-tailed resampling test) (Fig. 6D,E). The overall fraction of modulated neurons differed significantly between looped and non-looped neurons, independent of the threshold used (Fig S7I). Differences in the fraction excited or inhibited showed a consistent trend toward stronger modulation in looped neurons across all thresholds, although their significance depended on the distance threshold used to select non-looped neighbors of looped neurons (Fig. S7I). Thus, inputs from PM_V1_ neurons are more likely to modulate the visual responses of V1_PM_ feedforward-projecting neurons than those projecting to other targets, either through excitation or inhibition. Consistent with the enhancement of both inhibition and excitation of looped neurons during visual stimulation, photostimulation of FB axons did not result in a larger fraction of responsive looped neurons in the absence of visual stimulation (Fig. 6F).

We were unable to compare the extent of multiplicative and additive influences of FF and FB inputs in looped versus non-looped cells, as too few pairs of excited looped and non-looped cells with robust linear fits were located sufficiently close to each other to allow meaningful comparisons.

These results show that both FF and FB pathways interact differentially with neurons in the target area that project back to the source region, albeit in distinct ways. V1_PM_ inputs preferentially excite PM_V1_ neurons, whereas PM_V1_ neurons more selectively modulate V1_PM_ neurons both positively and negatively (Fig. 6G).

### Reciprocally connected populations show enhanced positive and negative correlations

We tested whether the preferential excitation of looped neurons by FF inputs, and their excitation or inhibition by FB inputs, is reflected in the structure of co-fluctuations measured in the multi-area recordings. As both FF and FB inputs show selectivity for reciprocating neurons, activity in V1_PM_ and PM_V1_ populations is expected to be preferentially coupled. However, the bidirectional recurrent selectivity for looped neurons precludes a clean isolation of the FF or FB effects on looped neurons using the FF_HIGH_ / FF_LOW_ or FB_HIGH_ / FB_LOW_ metric. Because FF is expected to preferentially excite FB-projecting neurons, and these in turn might selectively excite or inhibit the FF-projecting ones, the effects of FF and FB inputs on looped neurons are inherently entangled. Thus, because the activity of each looped population is both a cause and a consequence of the other’s activity, their respective contributions cannot be disentangled from the multi area recordings. However, this network organization is expected to result in stronger positive or negative correlations in V1_PM_ and PM_V1_ neurons relative to the general population. To test this, we measured noise correlations in pairs of V1-PM neurons and checked how they depended on projection type. A larger fraction of V1_PM_-PM_V1_ neuron pairs showed significant positive and negative correlations than V1_ND_-PM_ND_ pairs (Fig. 6H,I). Pairs including a reciprocating neuron and another one of unknown projection pattern showed intermediate values relative to the other groups. We conclude that the selective bidirectional recurrent engagement of reciprocally projecting populations revealed by optogenetic stimulation is reflected in the selectively enhanced shared variability of looped populations across areas, encompassing both positive and negative cofluctuations.

## Discussion

We used statistical analyses in multiarea recordings and optogenetic manipulations to dissect the influence of direct cortico-cortical projections on visual representations. We found that FF and FB projections had both inhibitory or excitatory influences and shaped feature tuning in their target area. Excitatory influences were more additive for FF than for FB inputs, whereas FB inputs exerted a comparatively stronger multiplicative influence on visual responses that boosted stimulus discriminability. The extent of inhibitory or excitatory influences depended on the projection type of the target neurons. FF inputs were more likely to excite reciprocating neurons, while FB inputs were more likely to either inhibit or excite in these neurons. This preference for recurrent interactions resulted in enhanced positive or negative co-fluctuations in neurons sending reciprocal projections.

The additive influence observed for FF inputs aligns with the established view that lower-order areas drive activity in higher-order ones; correspondingly, the multiplicative role of FB inputs is consistent with the classical notion that these projections modulate, rather than drive, activity in their targets. Our findings are also consistent with a role for corticocortical FB in mediating top-down modulation, such as during attention, and improving visual discrimination performance^15,16^. Our results show that these effects can be mediated by direct cortico-cortical inputs, without requiring indirect pathways such as cortico-thalamo-cortical or intermediate cortical areas. They also demonstrate that multiplicative FB effects are present even between early visual areas separated by a limited hierarchical distance, indicating that such modulation can emerge early in cortical processing. The stronger contribution of FB-driven multiplicative gain to stimulus decoding further supports the idea that the computational role of FB-mediated excitation is to refine the precision of cortical representations rather than to introduce new information.

The modulatory differences between FF and FB inputs are also consistent with their distinct dendritic distributions. Although both pathways innervate perisomatic and distal dendrites, FB inputs are enriched in the apical tufts, whereas FF inputs are more prevalent in perisomatic dendrites^5,6^. Our finding that FB pathways showing a stronger bias toward multiplicative effects and FF pathways toward additive effects is consistent with the idea that inputs to distal dendrites can exert multiplicative effects on somatic activity, whereas inputs targeting proximal dendrites more effectively contribute to action potential generation^52^. Although the balance of additive and multiplicative influences differed between pathways, neither was exclusive to one as both FF and FB projections gave rise to each type of effect across a substantial proportion of neurons. Our results therefore argue against a strict segregation of computational mode by projection direction, instead revealing a mixture of additive and multiplicative influences in both pathways, with graded biases rather than categorical distinctions. This is consistent with both pathways comprising driving and modulatory axonal connections^10,53^, albeit in different proportions in the FF and FB pathways.

Our observations also revealed that FF and FB influences depend on the projection type of their target neurons, with reciprocating neurons being preferentially excited or inhibited by these cortico-cortical inputs. These findings are consistent with theories of cortical computation in which cortico-cortical inputs preferentially engage reciprocating neurons to support predictive coding^17,18,54,55^ or to provide teaching signals during learning^56–58^. The different signs with which FF and FB inputs engage with looped neurons might reflect their different roles in these recurrent computations. FF inputs selectively excite FB-projecting, looped neurons, recruiting them in a way that closely links FB activity to the FF drive. In turn, FB inputs selectively excite or inhibit reciprocating, FF-projecting neurons. The selective modulation of FF-projecting neurons by FB inputs, with opposing signs, is well aligned with proposed circuit implementations of comparator circuits, in which descending predictions are compared with incoming sensory signals to generate positive and negative prediction errors^17^. Positive prediction errors arise when sensory inputs exceed expectations, whereas negative prediction errors occur when sensory inputs fall short of predicted signals. Proposed circuit implementations suggest that positive and negative prediction errors arise from differential actions of FB inputs on distinct L2/3 neuronal populations^59,60^. Specifically, FB inputs are thought to inhibit one population indirectly through the recruitment of local interneurons while directly exciting another population. Our results suggest that these positive and negative comparisons may be implemented selectively in FF-projecting neurons. These neurons, in turn, preferentially excite FB-projecting neurons, establishing a recurrent corticocortical circuit that may enable iterative comparisons across hierarchical levels. It remains an open question how the sign and magnitude of these cross-area influences depends on the tuning relationship of the lower-order and higher-order neurons, as predicted by the comparator framework.

The preferential excitation of reciprocating neurons observed for both the FF and FB pathways between V1 and PM may reflect their stronger innervation by these pathways^6,37,61^. However, the selective inhibition of reciprocating neurons would require recruitment of specific interneurons, as corticocortical pathways are almost exclusively excitatory. One possible circuit mechanism would rely on functionally organized interneuron pools, where local interneurons are wired to selectivity inhibit FF-projecting neurons according to their projection target. This would require target-specific wiring between FB afferents, local interneurons, and FF-projecting output neurons allowing inhibitory control over specific corticocortical pathways. However, selectively linking individual FB pathways to its reciprocating FF-projecting subpopulation would require a large variety of highly specialized pools of interneurons, with each pool comprising only a small number of neurons given the limited proportion of interneurons in cortical circuits. Thus, selective FB-mediated inhibition of reciprocating neurons might operate between functionally related group of areas such as dorsal or ventral visual stream areas^62,63^. Further work is needed to elucidate the circuit mechanisms underlying projection-type-specific inhibition by FB inputs.

Together, our findings reveal that direct cortico-cortical pathways can exert distinct and pathway-specific influences on sensory representations within recurrent cortical circuits. FF and FB inputs differentially combine additive multiplicative and inhibitory modulations with selectivity for reciprocating neurons, suggesting that direct cortico-cortical pathways are not simply broadcasting activity across areas, but are structured to reinforce specific recurrent circuits, These results highlight the functional specificity of direct cortical pathways and suggest that recurrent interactions between cortical areas may be a fundamental substrate through which sensory representations are dynamically shaped across the cortical hierarchy.

## Resource availability

### Lead contact

Requests for further information and resources should be directed to and will be fulfilled by the lead contact, Leopoldo Petreanu

### Materials availability

This study did not generate new, unique reagents.

### Data and code availability

Preprocessed two-photon data from the multi-area simultaneous recording and optogenetic stimulations will be deposited in Zenodo.

Code is published on GitHub: https://github.com/matthijsoudelohuis/vasile-oude-lohuis-et-al-2026-affinemodulation (analysis of the multiarea recordings), https://github.com/flora-vasile/vasile-oudelohuis-2026 (analysis of optogenetics experiments).

## Acknowledgements

We thank Ahmet K. Sarikaya and Emanuele Paoli for comments on the manuscript and the Champalimaud Research Hardware, Histopathology and Imaging Platforms for technical support. L.P. was supported by grants from Fundação para a Ciência e a Tecnologia (PTDC/MED-NEU/6645/2020), La “Caixa” Banking Foundation (HR22-00778) and by the Champalimaud Foundation. M.O.L. was supported by grants from the Dutch Research Council (NWO) under grant ID (019.223EN.013) and EMBO Postdoctoral Fellowship (ALTF 509-2023). F.M. was supported by a Transition To Independence grant from the Simons Foundation Collaboration on the Global Brain (00002313). C.K.M. was supported by Simons Foundation Collaboration on the Global Brain (543009 and 2794-0) and NIH (R01 EY035896 and RF1 NS127107).

## Author contributions

F.V., M.O.L. and L.P. conceived the study. F.V., D.P.S. and M.O.L performed the experiments. C.M. helped develop the optogenetic experimental methods and analyses. F.V, M.O.L. and F.M. analyzed the data. V.F., M.O.L. and L.P. wrote the paper. C.K.M. and L.P. supervised the project and secured funding.

## Methods

### Animals

Male mice (n = 8) of genotype GP4.3 (JAX#024275) were used in the multiarea recordings and male and female (n = 42) F1 offsprings from the Slc17a7-IRES2-Cre (JAX#023527) x Ai148D (Ai148(TIT2L-GC6f-ICL-tTA2)-D, JAX#030328) crossing mice were used for the optogenetic experiments. Both mouse lines expressed GCaMP6s heterozygously. Mice were group housed, maintained under a 12:12h regular light:dark cycle, at 23°C with a humidity of 45-65%, and had access to food and water *ad libidum*. All procedures were reviewed and approved by the Champalimaud Center for the Unknown Ethics Committee and performed in accordance with the Portuguese Direção Geral de Veterinária.

### Cranial window implantation and virus injection

Surgeries were performed on adult mice (8-16 weeks-old) under isoflurane anaesthesia (1.5-2%). Bupivacaine (0.05%; injected under the scalp) and buprenorphine (100 μl, 0.1 mg/kg, injected subcutaneously) provided local and general analgesia, dexamethasone (100 μl, 2 mg/kg, injected subcutaneously) and sodic cefovecin (100 μl/10 g, 6 mg/kg, subcutaneously) were used to minimize inflammation and infection, respectively. Eyes were protected and kept moist using ophthalmic ointment (Vitaminoftalmina A, Labesfal). While under anesthesia, body temperature was maintained at 37 °C using a heating pad (Supertech Instruments). Glass injection pipettes (Drummond) were beveled at 45° with a 13 µm inner diameter opening and backfilled with mineral oil and frontloaded with viral suspension before injection. A fitted plunger controlled by a hydraulic manipulator (Narashige, MO10) was inserted into the pipette and used to load and inject the viral solution. Per injection site, 25-30 nl was injected at two depths relative to dura (250-300 μm and 500-600 μm). To prevent backflow during withdrawal, the pipette was kept in the brain for 5 min following each injection. For the multiarea recordings, we targeted two virus injections to V1 at coordinates (posterior to Bregma 3.6 mm, lateral from midline suture 2.4 mm and posterior to Bregma 4 mm, lateral from midline suture 2.8 mm) and one injection to higher visual area PM (posterior to Bregma 2.7 mm, lateral from midline suture 1.6 mm). These coordinates were chosen based on overlap in retinotopic coverage between the areas^47^. In V1, a mixture of retroAAV-pgk-Cre (Addgene #24593) and AAV1-Ef1a-fDIO-tdTomato (Addgene #128434) was diluted with artificial CSF in a ratio 1.5 : 0.05 : 5. In PM, AAV5-CAG-Flex-tdTomato (Addgene #28306) and retroAAV-EF1a-Flpo (Addgene #55637) were mixed with aCSF in ratio 1:1:2. Through this viral strategy, neurons projecting to V1 cells expressed Cre recombinase, but only those in PM would express tdTomato. In a mirror fashion, all neurons projecting to PM express Flp recombinase, but only those in V1 expressed tdTomato. Note that by design, this viral strategy leads to retinotopically aligned feedforward and feedback labeled neurons. We observed no expression in two control animals in which only one of the two areas was injected. The virus to area pairing varied across animals and we observed similar expression profiles through Flp and Cre recombinase. For the optogenetic experiments, when the injection was in V1 (FF experiments), the skull was thinned above the posterior part of V1 and the glass injection pipette was lowered through the remaining bone down to the cortex. A 2:1 mix of AAV-Syn-ChrimsonR-tdT (Addgene #59171-AAV5) and either red fluorescent microspheres (Red Retrobeads IX, Lumafluor), pAAV-FLEX-tdTomato (Addgene #28306-AAVrg) or AAV-hSyn-DIO-mCherry (Addgene #50459-AAVrg) was injected at the following coordinates: posterior to Bregma, 3.8 mm; lateral to midline suture, 2.7 mm. For injection in PM (FB experiments), the same mix of virus was injected through the cranial window after localization of PM using intrinsic imaging. We used AAV5 for ChrimsonR expression, as this serotype results in limited retrograde infection in the neocortex^6,64^, minimizing opsin expression in looped neurons. A circular craniotomy was performed over the left visual cortex (multiarea recordings: before viral injection, 4 mm diameter, center coordinates, posterior to Bregma, 3 mm; lateral to midline suture, 2.1 mm; optogenetic experiments: after viral injection, 3 mm diameter, center coordinates, V1: posterior to Bregma, 3.5 mm; lateral to midline suture, 2.6 mm; PM: posterior to Bregma, 2.9 mm; lateral to midline suture, 1.8 mm). The dura was left intact. The imaging window was constructed from 3 layers of microscope cover glass (Fisher Scientific, no. 1) joined with an UV-curable optical glue (Norland). The window was embedded into the craniotomy and secured in place using black dental cement (Lang dental). A custom-designed stainless steel headpost was attached to the skull with cyanoacrylate glue and black dental cement to allow head-fixation during two-photon recordings. Mice were allowed to recover for three days, after which habituation to the experimental setup began.

### Intrinsic signal imaging

One to two weeks after surgery and before starting two-photon imaging, intrinsic signal imaging was used to obtain a retinotopic map of the primary visual cortex and surrounding higher visual areas, as described before^48^. Mice were head-fixed and lightly anesthetized with isoflurane (0.8-1.2%) and injected intramuscularly with chlorprothixene (1 mg/kg). The eyes were coated with a thin layer of silicone oil (Sigma-Aldrich) to ensure optical clarity during visual stimulation. Optical images of cortical intrinsic signals were recorded using a Retiga QIClick camera (QImaging) controlled using Ephus^65^ with a high magnification zoom lens (Thorlabs) focused on the brain surface under the glass window at 5 Hz. To measure intrinsic hemodynamic responses, the surface of the cortex was illuminated with a 620-nm red LED while drifting bar stimuli were presented to the right eye in a monitor. An image of the cortical vasculature under the window was obtained using a 535-nm green LED. Azimuth and elevation maps of the visual cortical areas were obtained by calculating the phase for each pixel of the discrete Fourier transform at the visual stimulation frequency, smoothed with a Gaussian filter (25μm s.d.) and visual field sign maps were computed as the sine of the angle between the gradients of the average horizontal and vertical retinotopic maps^48,66^. The hemodynamic delay was cancelled by subtracting the phase maps of the experiments from opposing moving stimuli.

### Head-fixed setup

#### Multiarea recordings

Mice were head-fixed and placed on a 3D printed circular running wheel (10 cm diameter). Running speed was recorded using a rotary encoder (MAE3, US Digital) at 1000 Hz. Mice were acclimated to head-fixation and spontaneous running before data collection. Infrared LEDs (850 nm) were pointed at the face of the mouse to enable infrared video acquisition. The videos were acquired at 10, 15 or 30 Hz using a camera (FLIR FL3-U3-32S2M-CS) with a zoom lens and an infrared filter (850 nm, 50 nm cutoff) to avoid detecting the two-photon laser (920 nm). Facial imaging data was processed using FaceMap^67^. Facial motion energy was obtained as the average pixel-wise intensity difference across time of an ROI covering the face. Mice were surrounded by three iPad retina displays (LG LP097QX1, 9.7 inch diagonal, 15x20 cm, 2048x1536 resolution, 60 Hz refresh rate) positioned at 10 cm distance from the head arranged at 90° angles to the left, front, and right of the animal, so that the animal’s head was approximately in the geometric center of the setup. The displays subtended 66.9° in elevation and 270° in azimuth. To prevent light contamination from the displays, the brightness was controlled using custom hardware based on the resonance scanner position: displays were off before (-2 μs) until after (+2 μs) each line scan, effectively illuminating the displays only during the scan turnaround time.

#### Optogenetics

Mice were placed in an acrylic tube and head-fixed. Visual stimuli were presented on a LED monitor (BenQXL2411-B 144 Hz 24 ”, stimulus presented at 60 Hz refresh rate) positioned 11 cm from the eye and at 30° relative to the axis of the body of the mouse to provide stimulation to the right eye. To prevent light contamination from the display we synchronized the stimulus presentation to the flyback of the resonant scanner using a custom electronic board and modifying the computer monitor to control its LED light source.

### Two-photon imaging

#### Multiarea recordings

Two photon imaging data were collected using a large field of view two-photon microscope^38^. The excitation wavelength was 920 nm (Alcor 920-2 XSight, SPARK Lasers). Laser power was set to exponentially increase with imaging depth with an exponent of 225 μm and typically ranged from 50 to 120 mW from superficial to deep imaging planes. The microscope was controlled by ScanImage 2016 (Vidrio Technologies). After the fluorescence light passed through the primary dichroic (Longpass 750nm dichroic, Alluxa) into the detector arm of the microscope, a second dichroic mirror (562 nm edge BrightLine, SemRock) divided the fluorescence path into two signals (GCaMP and tdTomato), each directed into one of two GaAsP photomultiplier tubes (PMTs) (Hamamatsu H11706P-40). The setup was assembled on a vertically mounted breadboard whose XYZ positions and rotation were controlled electronically via a gantry system (Thorlabs). Thus, to position the imaging objective with regards to the mouse, the position and rotation of the entire microscope were adjusted while the position of the mouse remained fixed. The headbar and cranial window were implanted over the left occipital cortex, with a slight tilt relative to horizontal. The mouse was positioned such that its head was horizontal level in the setup, while the rotation of the gantry was controlled to get the imaging objective horizontally level with the cranial window. This horizontal tilt was typically 15-19°. The entrance of the objective was light-shielded using tape, and immersed in water. The tape also prevented water evaporation during the experiment. For each mouse imaged, we typically spent the first imaging day optimizing the gantry and head fixation configuration (pitch and roll orientation) for planar imaging of the entire window.

#### Optogenetics

Two-photon imaging was performed from 2 to 12 weeks after cranial window implantation. We used a custom microscope (based on the MIMMS design, Janelia Research Campus, https://www.janelia.org/open-science/mimms) equipped with a resonant scanner, GaAsP photomultiplier tubes (10770PB-40, Hamamatsu) and a 16× (0.8 NA) objective (Nikon), controlled by ScanImage^68^ (Vidrio Technologies). O-rings were glued to the headpost, concentric with the center of the cranial window, to form a well for imaging, and filled with water-based ultrasound gel. To prevent stray light from the LEDs from entering the imaging well, a flexible, conical light shield was attached to the headpost and secured around the imaging objective. GCaMP was excited using a Ti:sapphire laser (Chameleon Ultra II, Coherent) tuned to 920 nm. We used GaAsP photomultiplier tubes (10770PB-40, Hamamatsu) and a 16x (0.8 NA) objective (Nikon). Laser power measured after the objective was between 26 and 40 mW. The objective was tilted such that its optical axis was perpendicularly aligned to the imaging window.

### Image acquisition

#### Multiarea recordings

We targeted four distinct regions in the left cortical hemisphere: primary visual cortex (V1), posteromedial cortex (PM), anterolateral (AL), and retrosplenial cortex (RSP). RSP data was not analyzed in this study. V1 and PM imaging regions were selected based on expression site. AL and RSP were targeted based on the intrinsic imaging maps. We acquired images across 8 planes at 5.36 Hz per plane at a resolution of 512 x 512 pixels (600 μm x 600 μm). For 9/12 recording sessions V1 and PM were imaged with 4 planes in each area (minimally spaced 50 μm in depth). For 3/12 recordings, V1 and PM were imaged with 3 planes each, and AL and RSP with 1 plane each. After the first imaging session, we took an image of the entire window, as well as a depth profile of the tdTomato expression. The window overview was obtained by averaging 100 frames of a 4 x 4 mm field of view at approximately 100-300 μm below dura. We obtained a Z-stack of the V1 and PM expression sites at 10 μm spacing from 0 to 750 μm below dura averaging 100 frames per depth (Fig. 1C).

#### Optogenetics

For FF experiments, PM was imaged and for FB experiments, V1 was imaged. When possible, several sessions were acquired per mouse, in different XY coordinates and or/different depths. Single-plane images were collected at ∼15 Hz, 1024x1024 (600 μm x 600 μm) pixels at a depth between 90 and 200 μm. After the last imaging session of each mouse, we acquired two Z-stacks with the laser tuned to 800 nm and 1000nm to obtain mCherry and tdTomato fluorescence profiles, respectively (0-300 μm, 5 μm step). Because the two-photon excitation spectrum of mCherry includes both 800 nm and 1000 nm, whereas tdTomato is selectively excited at 1000 nm, comparing the Z-stacks acquired at different wavelengths allowed us to confirm the presence of mCherry expressing retrogradely labeled cells and ChrimsonR-tdTomato expressing axons (Fig. 3B).

### Visual stimuli

#### Multiarea recordings

Stimuli were full-field drifting sinusoidal gratings (0.04 c.p.d, 1 Hz) drifting in the direction perpendicular to the orientation of the grating (8 orientations, 16 directions) with 200 repetitions each (n = 3200 trials). Stimuli were presented for 0.75 s with 1.25 s intertrial interval. Visual stimulus presentation was controlled using BONSAI and Bonvision^69,70^. We recorded behavioral and neural activity while mice spontaneously ran on the running wheel.

#### Optogenetics

Full-field sinusoidal gratings drifting in the direction perpendicular to the orientation of the grating were presented (4 orientations, 8 directions) with 20 repetitions each. Gratings presented had two of the following pairs of spatial and temporal frequencies: 0.02 c.p.d./0.5 Hz, 0.04 c.p.d./1 Hz, 0.16 c.p.d./0.5 Hz, 0.08 c.p.d./0.5 Hz, 0.16 c.p.d./1 Hz, 0.32 c.p.d./1 Hz. Stimuli were presented for 1 s with 4 s intertrial interval (grey screen). A recording session lasted 60 min. Visual stimuli were generated using Matlab and Pychophysics Toolbox^71^.

### Optogenetic stimulation

Axon photostimulation was achieved using a 633 nm one-photon laser (Cobolt 06-MLD) through the objective. Photostimulation consisted of 25 Hz pulses with a duration of 0.01 s and a peak power of ∼4mW after the objective. To prevent light contamination from the laser, we synchronized the photostimulation to the flyback of the resonant scanner. Photostimulation was triggered for 1 s during the visual stimulus presentation and in the absence of visual stimulus.

### Histology analyses

#### Histology

After completion of two-photon imaging experiments, mice were deeply anesthetized, transcardially perfused, and fixed overnight with 4% paraformaldehyde in 0.1 M phosphate buffer, pH 7.4. The brain was cut into 50-µm-thick coronal sections with a vibrating slicer (Leica). Slices were stained with 4′,6-diamidino-2-phenylindole dihydrochloride (DAPI) and imaged with the 20x objective of a Zeiss confocal 980 inverted microscope.

#### Spectral unmixing

To distinguish between mCherry and tdTomato in mice used for optogenetics, we used a ‘spectral unmixing’ approach using a Zeiss 980 inverted confocal microscope. Briefly, brain slices expressing either mCherry or tdTomato alone were excited at 561 nm wavelength to create a lambda stack of 8.8 nm wavelength band using a diffraction grating to spectrally separate the emitted light, which is then collected in a multichannel photomultiplier equipped with a linear array of 32 detectors. This allowed us to infer the spectral signature of each fluorophore from how signal level varied at the different emission bands. Then, these emission profiles were used in samples expressing both mCherry and tdTomato to determine the probabilistic contribution of one and the other fluorophore at each pixel in each emission band.

#### Laminar distribution of PM_V1_/V1_PM_ neurons and ChrimsonR axons

For quantification of the layer distribution of PM_V1_ and V1_PM_ neurons expressing mCherry, 5 animals and 3 slices per animal were used. Cells were detected using Cellpose^72^ within a 300 µm-wide column spanning the cortical depth area over 8 slices, 1.5 µm apart. The cortical depth was binned into 20 groups and the laminar distribution of looped cells was calculated as fraction over the cortical depth. For quantification of the layer distribution of PM_V1_ axons and V1_PM_ axons expressing ChrimsonR-tdTomato, 5 animals and 3 slices per animal were used. Vertical fluorescence profiles of tdTomato were measured using imageJ and normalized to the maximum fluorescence in each slice.

#### tdTomato axons fluorescence relative to PM_V1_/V1_PM_ neurons location

To quantify the FF/FB axon density relative to the position of PM_V1_/V1_PM_ neurons, we first identified PM_V1_/V1_PM_ mCherry+ neurons with Cellpose^72^ in coronal histological sections. Next, FF/FB axon density was quantified by measuring tdTomato+ axon fluorescence intensity in L1 and L2/3 (35% of the full depth of the cortex, from the pia). A fluorescence profile parallel to the pia was calculated, and background fluorescence was subtracted. The tdTomato fluorescence was aligned to the position of each mCherry+ neuron that was within 150 μm of the peak tdTomato fluorescence (which corresponds to an estimation of the field of view imaged with the 2-photon microscope).

#### Confirmation of the absence of retrograde expression of ChrimsonR-tdTomato

Retrograde expression of ChrimsonR-tdTomato would induce a confound in our evaluation of the effect of FF and FB pathways activation on reciprocating neurons. Indeed, by definition, retrograde expression would be biased to looped neurons and optogenetic activation would stimulate them directly rather than the axons innervating them. To assess ChrimsonR-tdTomato retrograde expression, we quantified the number of cells that were both expressing ChrimsonR-tdTomato and the retro-AAV used for identification of reciprocating neurons. To do so, we injected 2:1 mix of AAV-Syn-ChrimsonR-tdTomato (Addgene #59171-AAV5) and AAV-hSyn-EGFP (Addgene #50465-AAVrg) in V1 (AP: 3.8 mm, ML: 2.7 mm) or PM (through the cranial window after localisation of PM using intrinsic imaging). Using the GFP retro-AAV instead of mCherry retro-AAV allowed us to circumvent the complication of the proximity of tdTomato and mCherry excitation spectra. 2-3 weeks after injection, mice were perfused as described above, cryostat-sectioned in 20-μm-thick coronal slices. Coronal sections were stained with a rabbit anti-DsRed polyclonal antibody (<u>1:1000 dilution,</u> Takara, cat. #632496) and secondary AlexaFluor 555-conjugated secondary antibody (1:1000 dilution, ThermoFischer, cat. #A-21428) to amplify the ChrimsonR-tdTomato signal and then imaged with the 20x objective of a Zeiss confocal 980 inverted microscope. We quantified the fraction of looped neurons that also expressed ChrimsonR-tdTomato. We found retrograde ChrimsonR expression to be almost completely absent in looped cells in both V1 or PM. Injections in V1 lead to 2 out of 543 GFP+ cells in PM also expressing tdTomato (20 slices from 3 mice). Injections in PM resulted in 2 out of 846 GFP+ cells in V1 also expressing tdTomato (21 slices from 2 mice).

### Channel crosstalk correction in multiarea recordings

The dichroic mirror used to separate GCaMP and tdTomato fluorescence did not achieve full spectral separation, causing tdTomato fluorescence to bleed into the GCaMP signal. Although the tdTomato signal was not activity-dependent, we corrected for this bleedthrough prior to GCaMP ROI detection. Because the bleedthrough coefficient depended on PMT gain settings and recording depth, we estimated it separately for each imaging plane using robust regression of the red channel onto the green channel across a random sample of 100 frames. The GCaMP fluorescence was then corrected by subtracting the tdTomato signal, scaled by the estimated coefficient (the slope of the regression).

### Preprocessing of calcium data

Registration and regions of interest (ROIs) detection were performed using Suite2p^73^ with default settings for GCaMP6s. The output was manually curated to keep only somatic ROIs and characteristic calcium transients. Mean ROI signal was neuropil corrected using F = F − 0.7 × Fneu, where Fneu is the neuropil’s fluorescence. We used the spike deconvolution data from suite2p, which is based on the OASIS algorithm^74^. The deconvolved activity was divided by the baseline fluoresence F_0_ (calculated as the moving average over 60 s of baselines – optogenetics; or mean fluorescence across the session – multiarea recordings) resulting in the main activity metric events/F_0_ used throughout the analyses. For the multiarea recordings, a standardized noise level was computed for each ROI^75^ which captures the magnitude of frame-to-frame fluctuations in ΔF/F relative to shot-noise-driven fluctuations independent of frame rate:

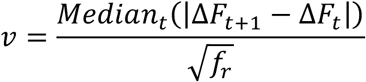

where ΔF is the delta F/F_0_ (prior to deconvolution), *t* is the frame, and *f_r_* is the frame rate. Neurons with noise level above 20 were excluded from all further analyses (4.777/51.709 ROIs, 9.2%).

### Labeled cell detection

#### Multiarea recordings

tdTomato expressing cells were based on automatic segmentation using Cellpose^72^. For each plane, the average motion-corrected tdTomato signal was square root transformed to accommodate the range from weak to strong tdTomato expression across cells. We trained a custom Cellpose model from on a subset of 20 planes with supervised labels. This custom model was subsequently applied to all planes and the output was minimally manually curated to remove labelled apical dendrites and add ROIS for missed cells. Projection neurons were identified based on a spatial overlap of the GCaMP ROIs (Suite2p output) and tdTomato ROIs (Cellpose output). Suite2p ROIs were routinely larger than cellpose ROIs as they contained fluorescence signal on dendritic and axonal segments. GCaMP ROIs were considered labelled cells if they contained more than 50% of the spatial footprint of a tdTomato ROI.

#### Optogenetics

After each recording session, a stack of 10 planes, 2 μm apart was acquired using Ti:sapphire laser (Chameleon Ultra II, Coherent) tuned to 830 nm for the red retrobeads or 1000nm for retro-mCherry and retro-tdTomato. Both GCaMP and retrobeads/mCherry/tdTomato were simultaneously acquired. To identify V1_PM_ and PM_V1_ neurons tagged with retrobeads/mCherry/tdTomato, we registered the GCaMP images from the stack acquired post-imaging session with the GCaMP images from the functional imaging using Fourier-based image correlation. Each plane of the stack was registered to the target functional imaging GCaMP and the one with the least error was kept for further processing. The corresponding retrobeads/mCherry/tdTomato plane was then overlayed to the ROI masks obtained from Suite2p in ImageJ and retrobeads/mCherry/tdTomato-positive cells were manually selected.

### Session inclusion criteria

For multi-area recordings, GCaMP and tdTomato maps were aligned to the intrinsic field sign maps primarily based on vasculature and used to establish the spatial extent of tdTomato expression (Fig. 1B). For optogenetic experiments, AAV injection target was verified post-hoc using QuickNII and VisuaAlign^76^ to align histology sections to the Allen Reference Atlas^77^. Recorded sessions from mice in which the injection location was outside of V1 for the feedforward experiments and outside of PM for the feedback experiments were discarded. For PM recordings, since the field of view recorded was larger than the PM area, we subselected PM neurons using the retinotopic map as a reference. Recording sessions were acquired at a minimum distance of 600 µm from the injection site to capture the effect of long-range interareal projections and avoid the influence of horizontal connections from neurons located close to the V1-PM border. To ensure successful optogenetic perturbation, only sessions in which 5% of neurons displayed significant positive or negative modulation by the photostimulation were included.

### Statistical Methods and Analyses

#### Feedforward and feedback population activity metric (multiarea recordings)

For each trial we computed a summary activity metric for the labeled projection neuron population, intended to capture the relative engagement of projection neurons versus local neurons in each area. The metric was defined as the difference between the mean activity across labeled neurons (V1_PM_ or PM_V1_) and the mean activity across nearby unlabeled neurons in the same area (V1_ND_ or PM_ND_) that were within 50 μm of any labeled neuron. Only sessions with at least 10 labeled neurons were included in the analyses to prevent estimating across area influences from few neurons. To characterize how neurons in the target area are modulated by the activity state of the projection population, trials were split into the bottom and top 25th percentile of this population activity metric, resulting into “low” and “high” conditions, respectively (FF_HIGH_ and FF_LOW_ for V1_PM_ – V1_ND_ and FB_HIGH_ and FB_LOW_ for PM_V1_ – PM_ND_). This was done per stimulus to ensure that any difference in neural responses between low and high conditions reflects modulation by the projection population state rather than differential stimulus selectivity between conditions. To minimize confounds of behavioral state, only trials with an average running speed below 0.5 cm/s and normalized facial motion energy (normalized 0–1 across the session) below 0.2 were included. This excluded 27.0% of trials on average across animals, although the proportion of spontaneous locomotion and facial motion varied across animals.

#### Computing overall modulation with discriminability index *d’* (multiarea recordings)

The target population were unlabeled neurons in L2/3 of the target area (recorded up to a depth of 270 μm). To assess whether a given target neuron’s overall response distribution was significantly shifted between the low and high activity conditions, a sensitivity index (*d’*) was computed. The low-condition and high-condition trial indices were pooled across all directions (preserving the per-direction split), yielding two large trial sets. The *d’* was defined as:

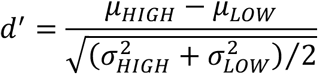

where μ and σ denote the mean and standard deviation of single-trial responses across all stimuli combined. To assess statistical significance, a permutation test with 1000 bootstrap iterations was used. On each iteration, for each direction, two subsets of 25% of the trials were drawn randomly from all still trials of that orientation (without reference to the population activity metric), and *d’* was recomputed on these random splits. A neuron was classified as significantly positively modulated if its observed *d’* exceeded the 99.5th percentile of the shuffle distribution (two-tailed α = 0.01), and as significantly negatively modulated if it fell below the 0.5th percentile.

#### Quantifying the effect of axonal photostimulation

For the optogenetics experiments, we first selected the responsive neurons by performing a paired t-test between the baseline (0-1 s of the trial) and stimulus (1-2 s of the trial) periods, on all visual stimuli and laser OFF and laser ON trials pooled (two-tailed, α = 0.01). We further classified these responsive cells as positively and negatively responsive by performing a t-test between laser OFF and laser ON trials (two-tailed, α = 0.01). Neurons for which the laser ON response was significantly different from and larger/smaller than the laser OFF response were considered as positively/negatively modulated by photostimulation. In the multiarea recordings, categorizing neurons as positively or negatively modulated neurons using the same statistical methods led to results that were 92% similar to the *d’*-based method described above.

#### Affine model

For each target neuron, the mean response across stimuli was computed separately for trials with and without photostimulation (for the axonal optogenetics) or low and high-activity trials based on the difference metric of labeled neurons (for the multiarea recordings). Mean responses were baseline subtracted to ensure response amplitude modulation was measured and baseline activity does not affect the modulation estimate.

Only cells with sufficient dynamic range in response to visual stimuli were included: at least a difference in mean evoked response of 0.04 (events/F_0_) between the minimum and maximum evoked response in control conditions (optogenetic experiments, N = 66/280 (FF) and 95/480 (FB) cells) or any of the low vs high conditions (multiarea recordings, N = 3016/5708 (FF) and 5307/8935 (FB) cells). This criterion was because multiplicative modulation cannot be assessed without variability in stimulus drive. Our results were similar when selecting neurons based on minimal orientation selectivity (gOSI > 0.4).

Stimulus conditions where the mean response was zero during photostimulation (optogenetics dataset) and FF_HIGH_/FB_HIGH_ (multiarea recordings), were treated as missing (set to NaN) to exclude saturating inhibition. Non-zero responsiveness was assessed by comparing the neuronal response between the baseline (0-1s of the trial) and the stimulus (1-2s of the trial) periods (paired t-test, α = 0.01). This was particularly important in the optogenetics dataset where some neurons were inhibited very strongly, creating a floor effect that precluded a clean estimate of subtractive versus divisive inhibition^78^. This saturating inhibitory effect was rare in the multiarea recordings, but for consistency we applied the same criterion across the datasets.

To characterize the nature of the cortico-cortical response modulation, the mean response across all stimulus conditions during high-activity (multiarea recordings) or laser ON (optogenetics dataset) was regressed against the responses during low-activity or laser OFF using ordinary least-squares linear regression. We used the responses to gratings at the two spatial frequencies we presented to fit the optogenetic measurements. To test for significance, a permutation test with 1000-5000 bootstrap iterations was performed. On each iteration and for each orientation, two random trial subsets were drawn without regard to photostimulation condition or the population activity difference metric, mean responses were computed and the regression was refitted. A neuron was classified as significantly multiplicatively/divisively modulated if its observed slope exceeded the 97.5th percentile or fell below the 2.5th percentile of the shuffle slope distribution (two-tailed α = 0.05). The same criterion was applied to the additive/subtractive offset.

#### Affine fit parameter control analyses

To confirm that differences in affine modulation parameters between the FF and FB pathways were not attributable to difference in response properties across the target area populations, the slope and offset parameters were analyzed as a function of gOSI (global orientation selectivity index), mean response amplitude, and the range of mean responses (across stimulus conditions) using analysis of covariance (ANCOVA). The effect of pathway on slope and offset parameters was tested when partialling out the contribution of each of these covariates and tested with the t-statistic for specific coefficients.

#### Von Mises tuning curve fitting

To characterize whether FF or FB modulation changed the preferred orientation or the tuning width of target neurons, a double von Mises function was fitted separately to the tuning curve under low/high conditions and photostimulation on/off. The double von Mises function was constrained to have two response components with stimulus directions 180° apart.

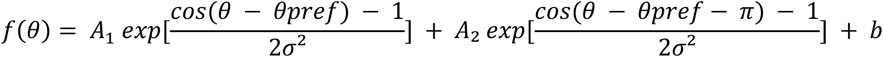

Where *θpref* is the preferred stimulus direction, *A_1_* is the amplitude of the response component at the preferred stimulus direction, *A_2_* the amplitude of the response component at the opposite stimulus direction, *b* is a constant baseline offset, and *σ* is the tuning width parameter where large values of *σ* correspond to broad tuning. Parameters were estimated by nonlinear least-squares optimization. Initial parameter estimates were derived from each neuron’s tuning curve, with the amplitude terms initialized from the response range and *θpref* from the orientation eliciting the maximum response, while parameters were bounded to physiologically plausible ranges (amplitudes between 0 and 1.5 times the response range, θpref within [0, 360 °], σ ≥ 7.5°, and b within [0, 1]). The difference in parameter values between the high and low conditions was compared to the difference obtained from two fits on two trial subsets obtained from a random selection of 25% of trials per stimulus direction.

#### Decoding stimulus features from population activity

To quantify the fidelity of sensory representations we decoded stimulus identity from small ensembles of simultaneously recorded neurons. We used a support vector machine (SVM) classifier trained with L2 regularization (λ = 0.5) and 5-fold crossvalidation. As a baseline characterization, we quantified how decoding performance varied across V1 and PM and scaled with population size using all available unlabeled L2/3 neurons and focusing on trials without photostimulation (optogenetics dataset) or without splitting trials into low- and high-activity pathway state (multiarea dataset). For each population size tested (N = 2-200), decoding was done n = 10 times with random subsets of neurons resampled without replacement and cross-validated decoding performance was averaged across resampled subsets to get one estimate per session.

To test whether corticocortical modulation affected the coding fidelity, decoding was performed separately for trials with and without photostimulation (optogenetics dataset) and trials split into low- and high-activity of difference metric of the labeled-unlabeled population in the other area (multiarea recordings). For each session, n = 10 neurons were randomly subsampled n = 100 times. We chose n = 10 neurons to (1) include also those sessions where fewer responsive neurons in the target area were simultaneously recorded and (2) relate the variability in multiplicative and additive variability of this randomly sampled small ensemble to decoding performance improvements (see below). Cross-validated decoding performance averaged across folds and resamples was compared between conditions and pathways with a repeated-measures ANOVA with session as the subject variable and activity level (low/high) and pathway (FF/FB) as within-subject factors. To assess the relationship between population coding fidelity and single-neuron gain modulation, the decoding performance ratio (high/low or on/off) was regressed against the mean slope and offset parameter values of the decoded ensemble^43^.

#### Assessing selectivity of photostimulation effect for reciprocally projecting neurons

Because the effect of photostimulation was anatomically organized, evaluating its specificity on reciprocating neurons required spatially local comparisons. This local comparison was necessary because the impact of photostimulation was spatially organized, the magnitude of laser-induced modulation being more similar between nearby neurons than between more distant pairs (Fig. S1). Moreover, because both FF and FB connections are retinotopically matched, FF and FB axons were more abundant around retrogradely labeled neurons (Fig. S7E,F), a relation that was particularly strong for FF axons and PM_V1_ labeled neurons. Consistent with this anatomical organization, unlabeled PM neurons located within 75 µm of a labeled neuron were more likely to be modulated by axonal photostimulation than neurons located farther away (Fig. S7G). Together, these observations indicate that comparisons between looped and non-looped neurons must be restricted to spatially neighboring cells to avoid confounding differences in local axonal density and photostimulation strength. To this end, we used a bootstrap resampling approach that controls for spatial variability. For each looped cell, we identified candidate non-looped neighbors from the same imaging session that fell within a distance threshold (ranging from 40 to 100 µm in 10 µm steps), computed as Euclidean distance between neurons. Looped neurons with fewer than two eligible neighbors within a given threshold were excluded from the analysis at that threshold. For each metric (fraction of excited, inhibited or modulated neurons) we computed the real mean across all retained looped cells. To generate a null distribution, we performed 10,000 bootstrap iterations: in each iteration, for every retained looped cell, one of its neighbors was randomly drawn, and the mean of the sampled metric was computed across all looped cells. Two-sided p-values were computed as twice the proportion of bootstrap samples above or below the real value.

#### Pairwise noise correlations

We computed noise correlations between pairs of neurons on the time-averaged response (0-1 s post stimulus) across trials of the same stimulus using a standard Pearson’s correlation coefficient. Noise correlation values were averaged across stimulus conditions to obtain one correlation value per neuron pair. The number of neuron pairs for each interarea population combination was: V1_ND_-PM_ND_ 66.168 ± 80.656, V1_ND_-PM_V1_ 13.222 ± 17.481, V1_PM_-PM_ND_ 9.889 ± 14.709, V1_PM_-PM_V1_ 1.970 ± 2.758. Significance of the pairwise correlations were tested by computing the noise correlations on trial-shuffled data, destroying the trail-by-trial correspondence between pairs of neurons but preserving the activity distribution. Pairwise correlations were deemed significant if they exceeded 99.995 percentile or fell below the 0.005^th^ percentile of this shuffled distribution (two-tailed α = 0.0001). To test whether the results depended on the distance to labeled neurons, the fraction of significantly correlated pairs was computed between pairs of neurons that were required to be within an increasing radius threshold to any labeled neuron. This was only performed if the two sets of neurons across areas consisted of at least 5 neurons in each population each to estimate of the fraction of significantly correlated pairs for that session and radius threshold.

### Statistics

Data in the text is mean±standard deviation. Statistical tests details can be found in the Methods, Results and figure legends.

---

**Figure S1.**
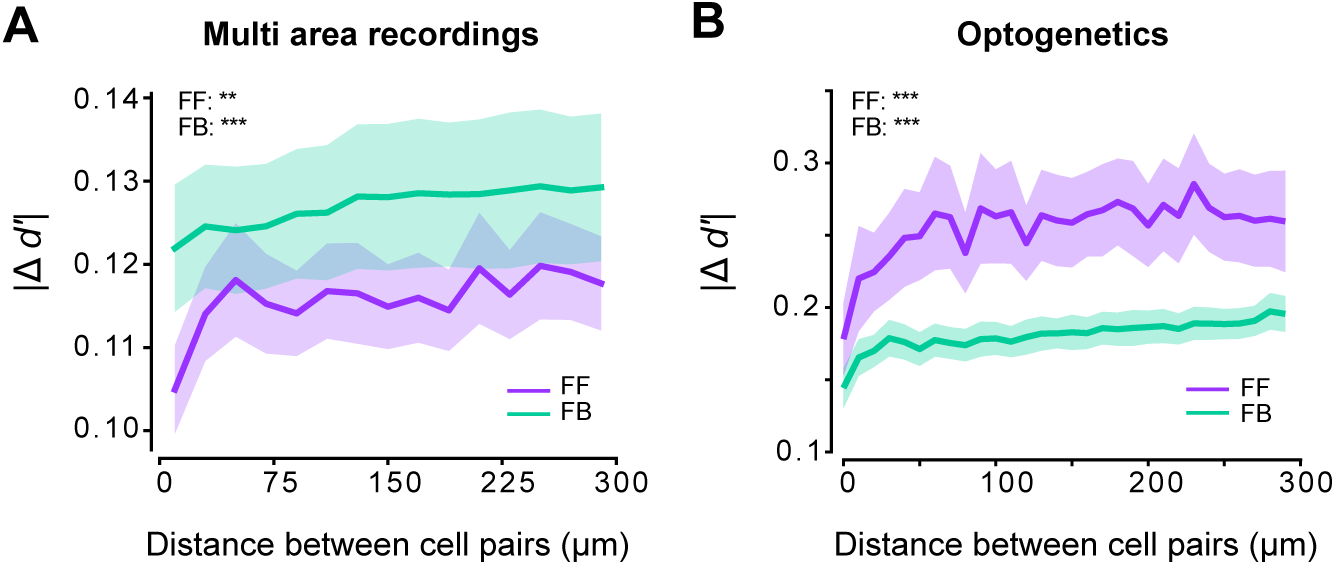
Neighbouring neurons are influenced by FF and FB more similarly. **A)** Quantification of the absolute difference in modulation between pairs of neurons as a function of the distance between them in the multi area recordings. Both high FF and high FB induce more similar modulation in neurons that are in close proximity (one-way repeated measures ANOVA per pathway, FF: F=60.9, p=1.28 x 10^-12^, FB: F=6.7, p=0.010. Line and shading indicate mean and s.e.m. across sessions (n=12 from N=8 mice). **B)** Same as a. for optogenetics experiments. Friedman test per pathway, FF: χ(29)=114.6, p=3.87 x 10^-12^, FB: χ(29)=180.2, p=9.09 x 10^-24^. Line and shading indicate mean and s.e.m. across sessions (FF: n=30 from N=16 mice, FB: n=23 from N=12 mice).

**Figure S2:**
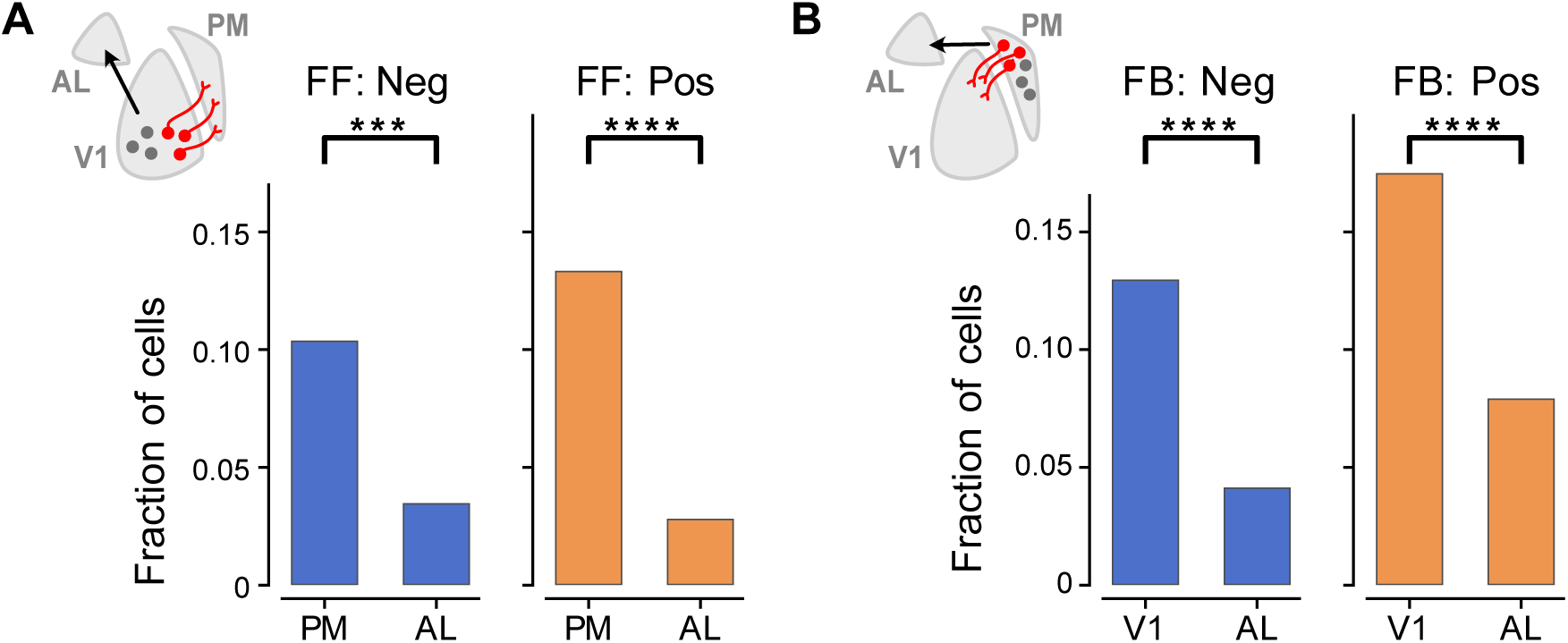
Projection neurons modulate activity in a projection-target specific manner. **A)** Fraction of negatively and positively modulated neurons by the FF metric (V1PM-V1ND) in visual areas AL and PM. A significantly smaller fraction of AL neurons than PM neurons was modulated by the state of the V1 to PM pathway (negative: ꭕ^2^ = 14.41, p = 0.00147, positive: ꭕ^2^ = 27.55, p = 1.53 x 10^-7^; n = 5943 PM neurons, N = 8 mice; n = 302 AL neurons, N = 3 mice). **B)** The same but testing the specificity of the labeled PM to V1 feedback pathway. The fraction of negatively and positively modulated neurons was lower in AL than in V1 (negative: ꭕ^2^ = 27.04, p = 1.99x10^-7^, positive: ꭕ^2^ = 24.76, p = 6.5x10-7; n = 10197 V1 cells, N = 8 mice; n = 410 AL cells, N = 3 mice).

**Figure S3.**
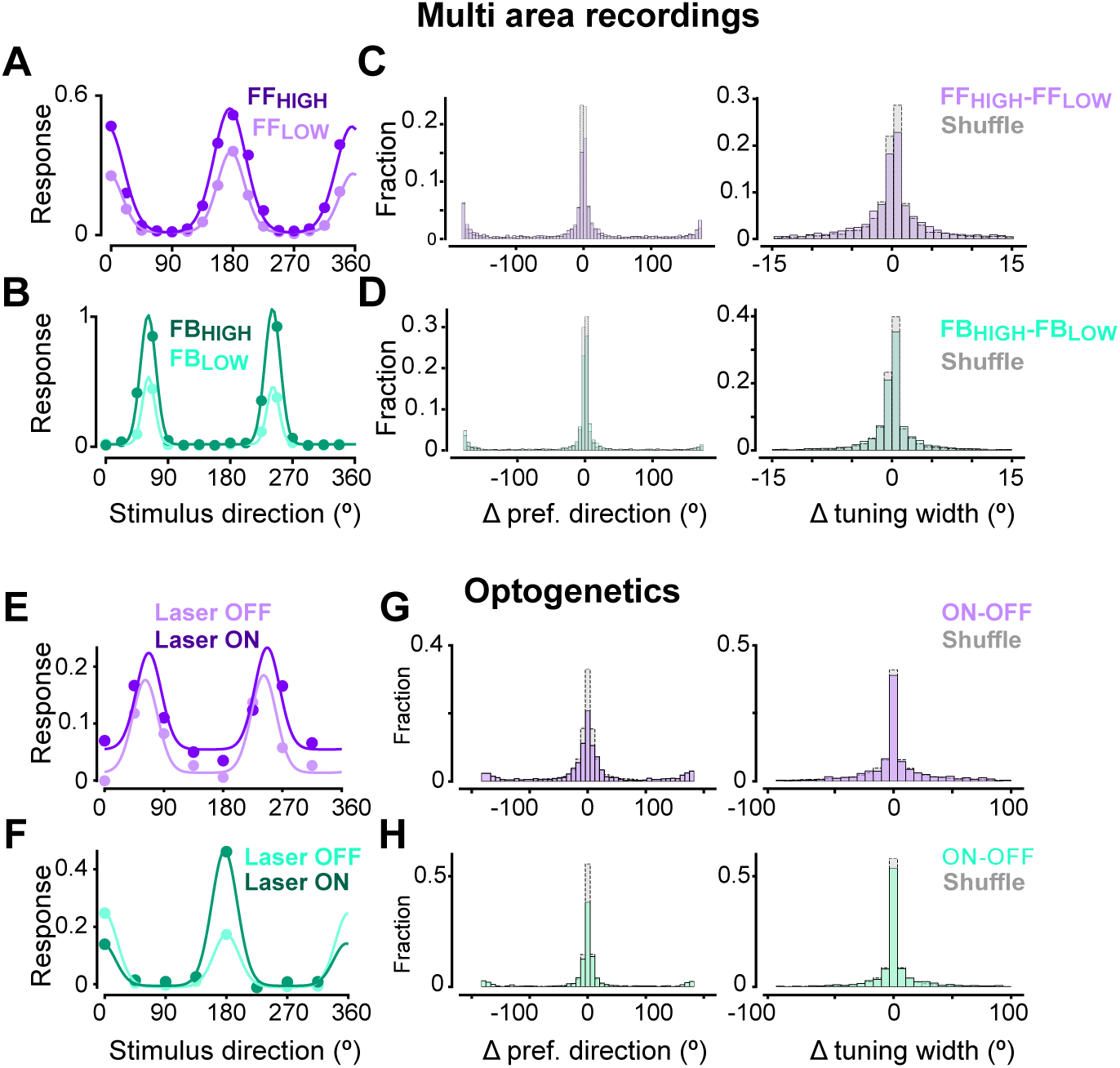
Preferred direction and tuning width remain unchanged across FF and FB HIGH/LOW trials and photostimulation. **A)** Example von Mises fit on a PM neuron during FF_LOW_ and FF_HIGH_ trials. **B)** Same as A, but for a V1 neuron during FB_HIGH_ and FB_LOW_ trials. **C)** Difference in preferred direction (left) and tuning width (right) of PM neurons between FF_LOW_ and FF_HIGH_ trials. Grey indicates shuffle with the difference in parameter values between two fits on random subsets of 25% of trials. The difference in tuning parameters is within the variability observed in random subsets of trials. n = 3016 neurons from N = 8 mice. **D)** Same as C, but difference for V1 neurons during FB_HIGH_ and FB_LOW_ trials. n = 5306 neurons from N = 8 mice. **E-H**): Same as A-D but for optogenetics experiments. Grey indicates shuffled Laser OFF and Laser ON trials. FF: n =1092 neurons from N=16 mice, FB: n =1381 neurons from N=12 mice.

**Figure S4.**
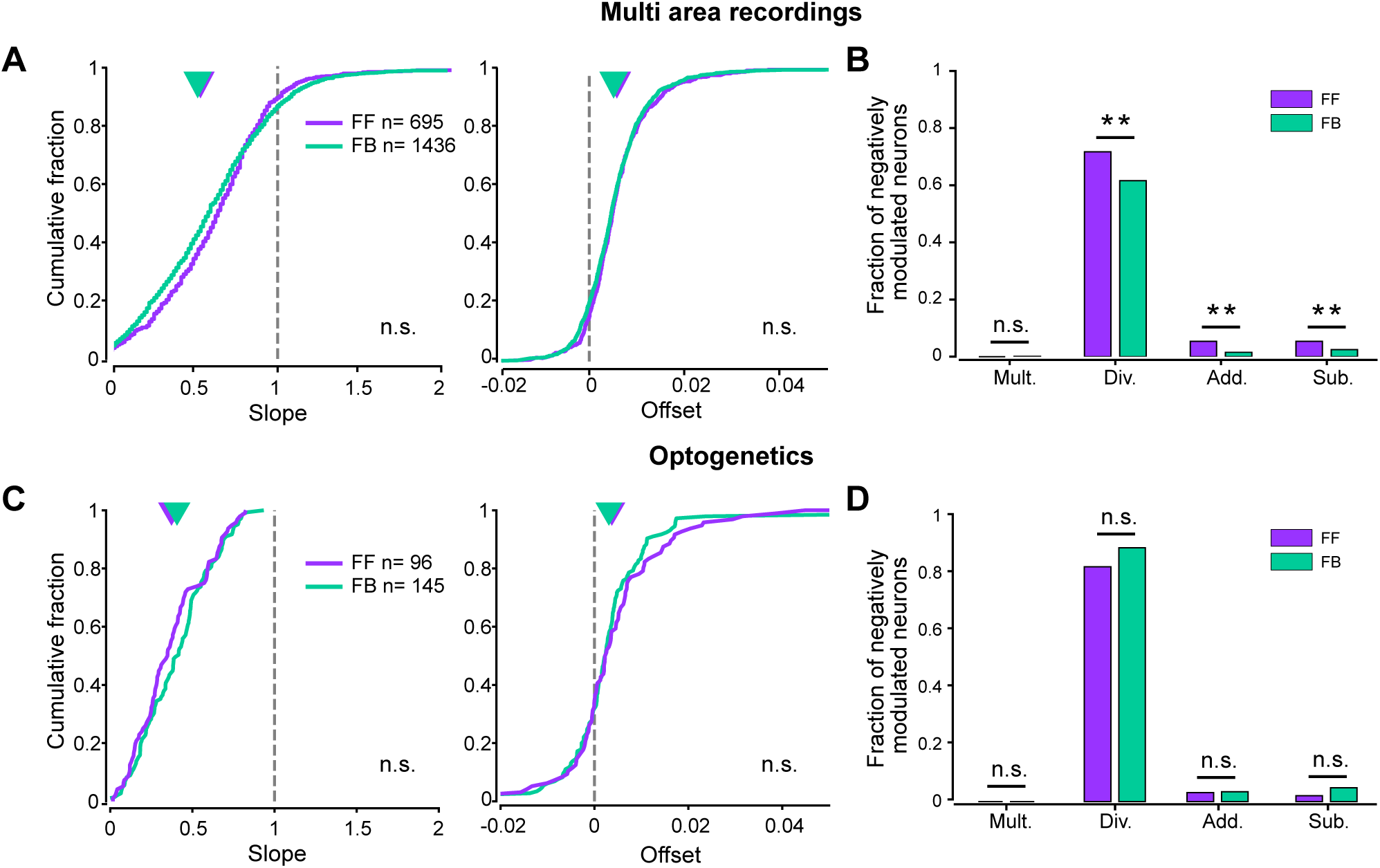
Effect of FF and FB pathways on the visual responses of negatively modulated cells. **A)** Cumulative distribution of multiplicative slope (left) and additive offset (right) of cells negatively modulated by high FF/FB activity. Slope: FF: 0.455, FB: 0.461, p=1; Offset: FF: 0.005, FB: 0.004, p=0.0912, Mann-Whitney U test with Bonferroni correction for multiple comparisons. N=8 mice. **B)** Fraction of significantly multiplicative, divisive, additive and subtractive neurons. Asterisks denote significance of a chi-square test of proportions (Div. χ = 11.28, p = 0.0063, Add. χ = 12.51, p = 0.0032, Sub. χ = 10.89, p = 0.0077). **C-D**): Same as A,B but for FF/FB photostimulation. For each neuron, only stimuli for which the visual response are not completely abolished are used for the fit. Slope: FF: 0.372, FB: 0.406, p=0.18; Offset: FF: 0.003, FB: 0.003, p=0.49, Mann-Whitney U test. FF: N=10 mice, FB: N=10 mice.

**Figure S5.**
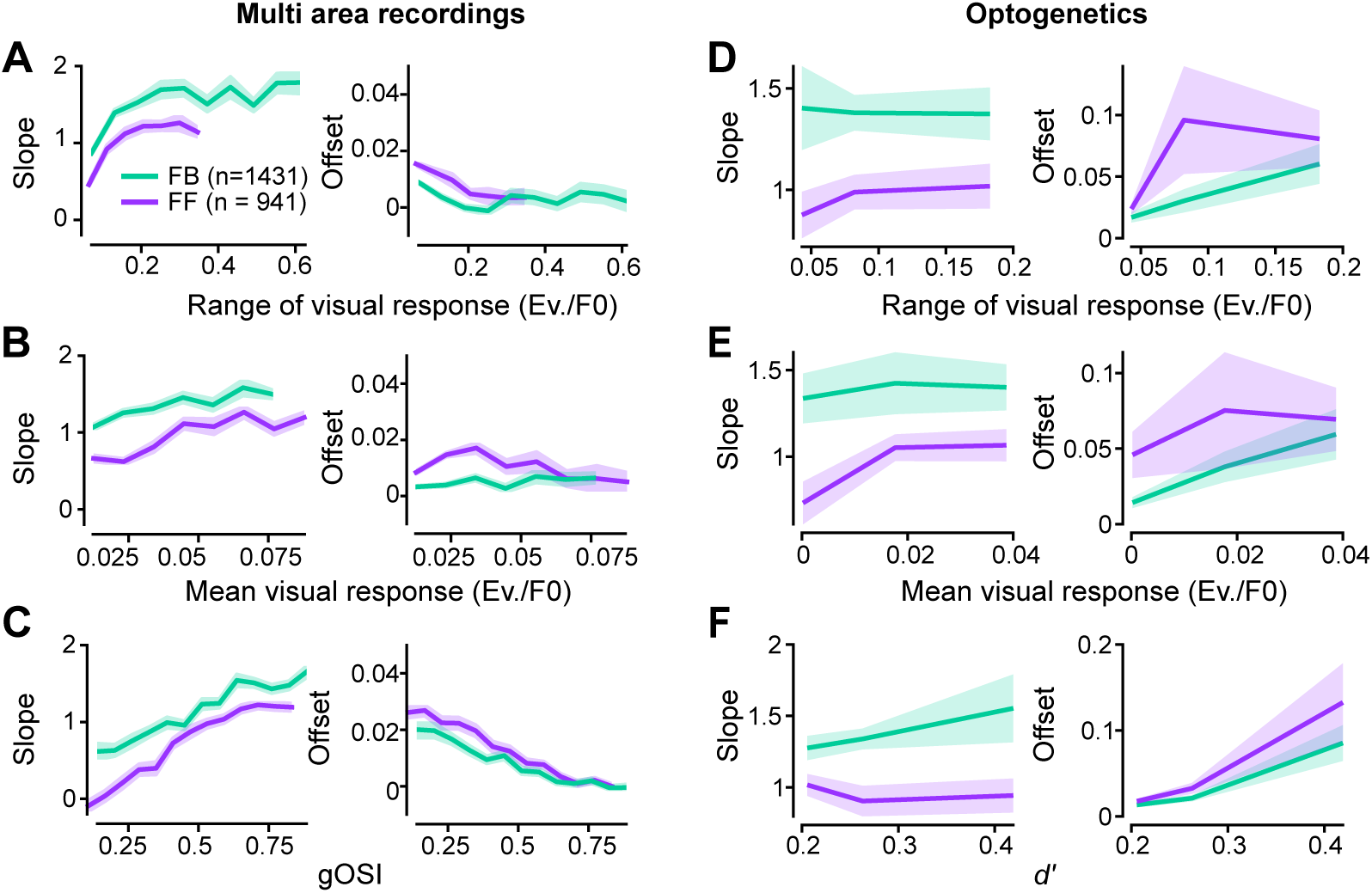
Pathway-specific slope and offset differences do not depend on visual response range and amplitude, orientation selectivity or *d’*. **A)** Slope and offset as a function of the range of the visual response across stimulus conditions for multi area recordings. t-statistic and p-value for the pathway effect (ANCOVA). Slope: t = 13.7, p = 1.74 x 10^-41^. Offset: t = -9.1, p = 1.57 x 10^-19^. N = 8 mice. **B)** Same as A for mean visual response. Slope: t = 15.9, p = 1.51 x 10^-54^. Offset: t = -10.0, p = 2.75 x 10^-23^. **C)** Same as A for global orientation selectivity index (gOSI). Slope: t = 11.6, p = 1.47 x 10^-30^. Offset: t = -5.6, p = 2.74 x 10^-8^. **D)** Slope and offset as a function of the range of the visual response across stimulus conditions for optogenetics experiments. Slope: t = -3.3, p =0.001. Offset: t = 2.1, p = 0.038. **E)** Same as D for mean visual response. Slope: t = -3.2, p =0.001. Offset: t = 1.8, p = 0.067. **F)** Same as D for *d’*. Slope: t = -3.5, p =0.0006. Offset: t = 0.9, p = 0.356.

**Figure S6:**
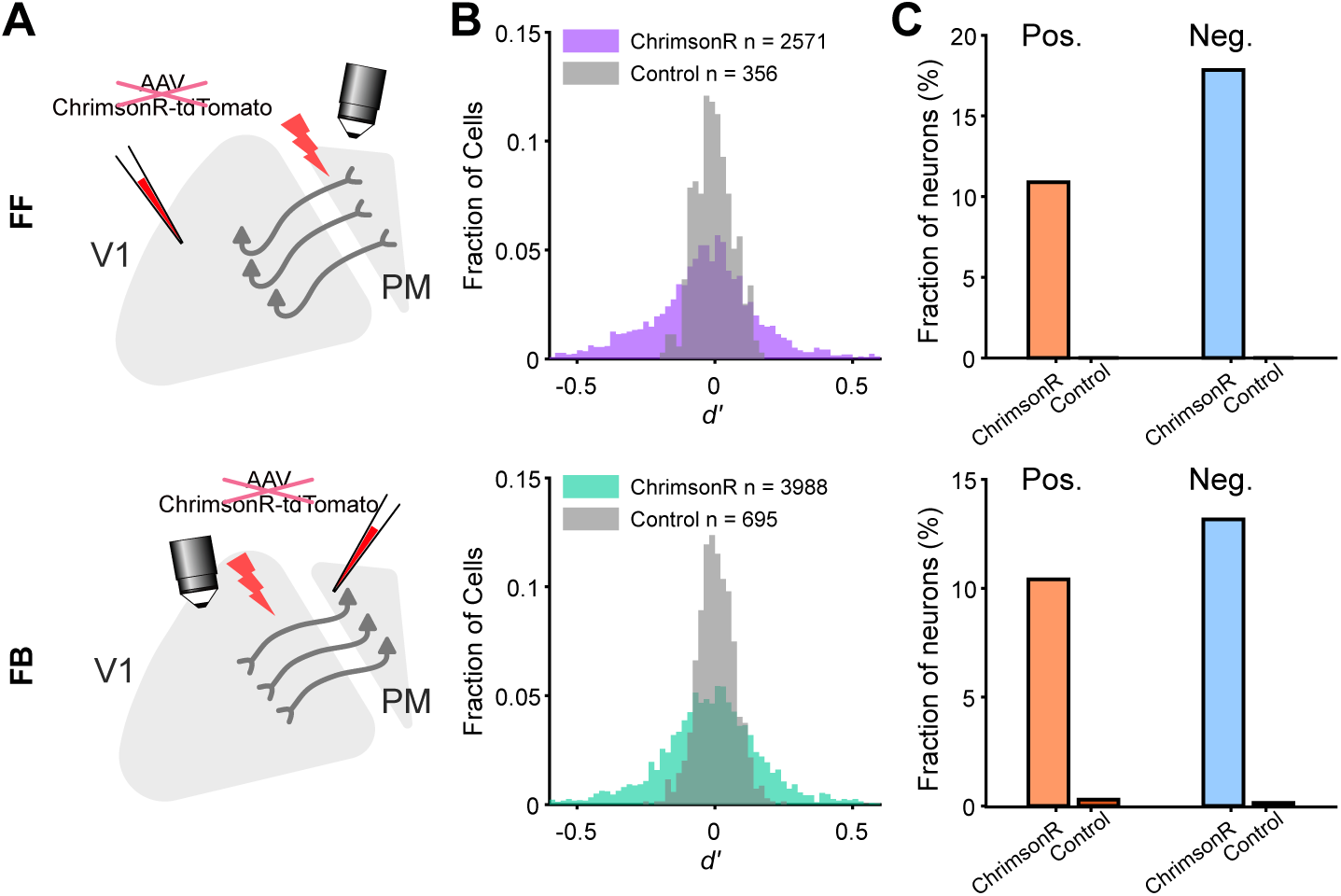
Photostimulation without ChrimsonR expression does not induce visual response modulation. **A)** Scheme depicting the experimental paradigm: photostimulation and 2P recording in PM (top) or V1 (bottom) without injection of the ChrimsonR-tdTomato AAV. **B)** Distribution of FF (top) and FB (bottom) photostimulation modulation in mice expressing (colored; FF: N=3 mice; FB: N=2 mice) or not expressing (gray) ChrimsonR. **C)** Fraction of significantly positively (orange) and negatively (blue) modulated neurons by FF (top) and FB (bottom) photostimulation with and without ChrimsonR.

**Figure S7:**
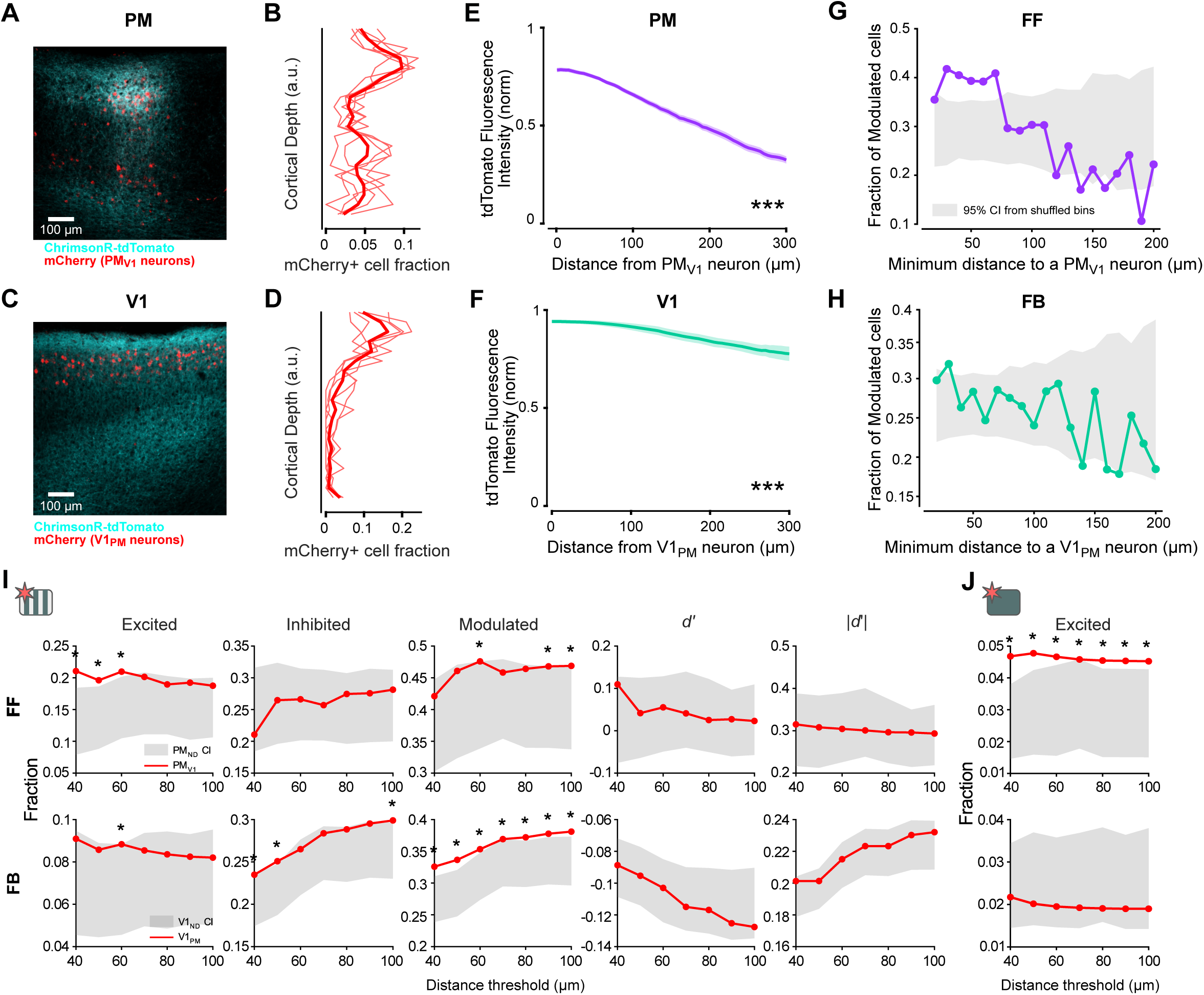
Spatial organisation of the influence of FF and FB photostimulation. **A)** Coronal section with V1PM ChrimsonR-tdTomato-expressing axons and PMV1 mCherry-expressing retrogradely labelled cells from the FF photostimulation experiments. **B)** Laminar distribution of PMV1 mCherry-expresing retrogradely labelled cells. Thick line, average, thin line, individual mice (n=5 mice). **C)** Coronal section with PMV1 ChrimsonR-tdTomato-expressing axons and V1PM mCherry retrogradely labelled cells from the FB photostimulation experiments. **D)** Laminar distribution of V1PM mCherry-expressing retrogradely labelled cells (n=5 mice). **E)** Fluorescence intensity of tdTomato-expressing FF axons as a function of the distance to mCherry-expressing PMV1 cells.Thick line, average; shading, s.e.m across 20 slices from N=5 mice (one-way repeated measures ANOVA, F_60,1140_ = 210.1, p = 3.02 x 10^-21^). **F)** Fluorescence intensity of tdTomato-expressing FB axons as a function of the distance to mCherry-expressing V1PM cells (one-way repeated measures ANOVA, F_60,480_=21.6, p= 9.98 x 10^-5^, n=9 slices, N=4 mice). **G)** Fraction of PM neurons modulated by FF photostimulation (excited or inhibited) as a function of their distance to a PMV1 neuron. Grey shaded area corresponds to the 95% confidence interval from shuffled bins. **H)** Same as G for the fraction PM neurons modulated by FB photostimulation and distance to a V1PM neuron. **I)** Same analyses as Fig 6B,C but for different distance thresholds for selecting non-reciprocating neurons in the vicinity of looped neurons. The fraction of excited, inhibited and modulated reciprocating neurons (red line) is compared with shuffled resamples of neighboring non-reciprocating neurons at different distance thresholds (grey distribution, 95% confidence interval from 10.000 resamples). Top row, FF photostimulation. Bottom row, FB photostimulation. **J)** Same as I for the fraction of excited cells by photostimulation without visual stimulus.

